# Micafungin exposure drives multidrug resistance in *Clavispora lusitaniae*

**DOI:** 10.64898/2026.06.29.735437

**Authors:** Elizabeth Wash, Nancy E. Scott, Katura Metzner, Xin Zhou, Deveney Da Silva, Nivea Pereira De Sa, Sana Akhtar Usmani, Nathalia Fidelis Viera De Sa, Maurizio Del Poeta, Anna Selmecki

## Abstract

Fungal infections are an escalating global health concern, with rare *Candida* species posing an urgent threat due to emerging multidrug resistance. *Clavispora* (*Candida*) *lusitaniae* is an uncommon pathogen in which multidrug resistance has been documented during antifungal therapy, yet the selective forces driving this phenotype remain unclear. Here, we show that exposure to the echinocandin micafungin (MCF) alone can select for multidrug resistance in *C. lusitaniae*. Through controlled evolution experiments we identified individual point mutations in genes encoding ergosterol biosynthesis enzymes (*ERG*s), sterol trafficking proteins (*OSH2*), and the echinocandin drug target (*FKS1*) that confer a significant fitness benefit to one or more classes of antifungals. We find that *ERG3* loss-of-function is the primary and independent driver of pan-antifungal resistance to echinocandins, azoles and polyenes. The *ERG3* mutants have <1% ergosterol, increased levels of non-toxic sterol intermediates, and increased chitin content, consistent with both cell membrane and cell wall remodeling that enables the fungal pathogen to evade all three drug classes. The convergence of sterol reprogramming and compensatory cell wall remodeling that occurs during adaptation to echinocandin monotherapy can evolve through a single point mutation and parallels our recent case study of acquired multidrug resistance.

**IMPORTANCE:** Multidrug resistance in *Candida* species severely limits treatment options and increases mortality, particularly in immunocompromised patients. Despite increasing reports of multidrug resistance, the molecular mechanisms driving multidrug resistance remain poorly understood. We find that *in vitro* MCF exposure alone can drive multidrug resistance in *C. lusitaniae* via acquisition of *de novo* point mutations in *ERG3,* an observation that parallels our recent patient case study. By identifying causative mutations and associated physiological changes, we provide mechanistic insight into the emergence of multidrug resistance and highlight the need for surveillance strategies that account for resistance evolution under echinocandin monotherapy.

## INTRODUCTION

*Candida* species are opportunistic fungal pathogens that can cause life-threatening bloodstream infections (candidemia). *C. albicans* is the most common etiological agent of candidemia, but non-*albicans Candida* are increasingly prevalent (1, 2). *Clavispora* (*Candida*) *lusitaniae* is an emerging haploid fungal pathogen similar to *Candidozyma* (*Candida*) *auris*, with a propensity to acquire antifungal resistance (3). Reports of candidemia cases caused by *C. lusitaniae* have increased ∼4-fold from 2014 to 2019 in the US (4), in part due to the rapid acquisition of antifungal drug resistance during antifungal therapy (5).

Only three major classes of antifungals (echinocandins, azoles, and polyenes) are used to treat candidiasis, and all target essential components of the fungal cell wall or membrane (6). Echinocandins, including micafungin (MCF), are fungicidal and the standard of care for invasive infections in the US (6). Echinocandins inhibit the essential catalytic subunit of 1,3-β-D-glucan synthase encoded by *FKS1* (7, 8). Non-competitive inhibition of Fks1 disrupts cell wall biosynthesis and leads to cell death (9–11). Azoles, including fluconazole (FLC), are fungistatic and a common step-down therapy following echinocandin treatment (12). Azoles interfere with the biosynthesis of ergosterol, a crucial component of the fungal plasma membrane, by directly inhibiting lanosterol 14-α-demethylase encoded by *ERG11* (13). Azoles perturb membrane stability and arrest cell growth due to the depletion of ergosterol and an accumulation of a toxic 14α-methyl-ergosta-8,24(28)-dien-3β,6α-diol sterol synthesized by Erg3 (14–17,18). Production of the toxic sterol is variable across fungal pathogens due to differences in Erg3 enzyme activities, which is considered an important mediator of azole susceptibility (19). The final drug class is the fungicidal polyenes, including amphotericin B (AMB), that bind ergosterol directly resulting in membrane destabilization, formation of membrane pores resulting in ion leakage, and accumulation of reactive oxygen species (20–23). Critically, both *C. lusitaniae* and *C. auris* have a propensity to develop resistance to AMB during in-patient therapy (24–27).

Resistance to all three major classes of antifungal drugs across *Candida* species creates treatment challenges (1, 26, 28, 29). Mutations that alter the binding of drug targets, like Fks1 and Erg11, are common in resistant isolates (30–33). *FKS1* mutations, primarily in hotspot domains, are the major driver of echinocandin resistance both *in vitro* and in clinical isolates (34–37). *FKS1* mutations can decrease drug binding or alter the enzyme’s catalytic capacity, leading to a decrease in 1,3-β-D-glucan abundance in the cell wall (30, 31), frequently with a compensatory increase of chitin synthase activity to support the compromised cell wall (38–40). Increased chitin synthesis is a conserved mechanism conferring reduced echinocandin susceptibility across *C. albicans*, *C. tropicalis*, *C. parapsilosis*, and *C. guilliermondii* isolates (41). Mutations outside of *FKS1* that drive decreased susceptibility to echinocandins are rare but include the sphingolipid biosynthesis gene *FEN1^Y298*^* in *Nakaseomyces glabratus* (*Candida glabrata*) (42).

Resistance to the azole class of antifungals can be conferred by changes to lipid synthesis (24, 43–48) and/or drug efflux (49, 50). A common mechanism that confers azole resistance is mutations in the drug target *ERG11* (32, 51, 52). However, drug target mutations do not always recapitulate resistance to antifungals across species. For example, cloning *ERG11^I466M^* or *ERG11^Y501H^* mutations identified in azole-resistant *C. auris* into *S. cerevisiae* did not confer azole resistance (53). Non-target mutations are increasingly recognized in azole-resistant isolates, indicating that alternate mechanisms contribute to azole resistance (54–56).

Mutations in non-target genes can confer multidrug resistance, i.e., resistance to two or more classes of antifungals. Reports of multidrug resistance in diverse *Candida* species, including *C. lusitaniae, C. parapsilosis, C. auris, N. glabratus, C. tropicalis, C. krusei,* and *C. blankii* are increasing in incidence and vary by geographic region (57–66). We recently reported a ∼2-fold increased rate of resistance to FLC, MCF and multidrug resistance in non-*albicans* species in the Twin Cities of Minnesota relative to other U.S. regions (1). Multidrug resistance to azoles and polyenes, which share ergosterol as a target, is repeatedly identified in clinical isolates (15, 67, 68). However, non-target mutations have also been identified in isolates with resistance to drugs that do not share targets. Mutations in *BLM3*, *CDC10,* and hypothetical proteins are associated with multidrug resistance to FLC and caspofungin in *C. auris*; *in N. glabratus IPI1* mutations confer resistance to MCF and multiple azoles; and mutations in *ERG3* in *C. lusitaniae* and *C. parapsilosis* confer multidrug resistance to azoles and echinocandins (64, 69–71). Despite the increasing prevalence of multidrug resistance, the mechanisms by which non-drug-target mutations confer multidrug resistance are largely uncharacterized and require molecular engineering to identify causality for each unique instance.

We recently reported the evolution of multidrug resistant *C. lusitaniae* from a patient treated with a single echinocandin (MCF), over an 11-day period (64). The 22 serial isolates from this patient were clonal and differed by only 0-3 unique single nucleotide polymorphisms. All initial isolates were pan-drug sensitive, but echinocandin resistance and echinocandin-azole multidrug resistance rapidly emerged, despite no prior history of azole therapy. Multidrug resistance correlated with the acquisition of a single, *de novo* missense mutation *ERG3^Q308K^*(64). *ERG3* encodes a C-5 sterol desaturase that converts C-5(6) saturated sterols (like episterol) to ergosta-5,7,24(28)-trienol (72), though in the presence of azole drugs, Erg3 synthesizes a toxic sterol diol that disrupts membrane function (16). Accordingly, Erg3 loss of function mutations cause azole resistance due to a lack of this toxic sterol, and polyene resistance due to the reduced production of ergosterol (18, 46, 67, 68, 73, 74). However, the mechanism by which *ERG3* mutations confer echinocandin resistance is unknown, and it is unclear whether MCF exposure is sufficient to select for multidrug resistance in *C. lusitaniae*. Furthermore, because *ERG3* loss of function mutations often co-occur with *FKS1* mutations in clinical isolates of diverse *Candida* species (*C. parapsilosis*, *N. glabratus*, *C. auris*, *C. albicans*, *C. lusitaniae*) (75), it is unclear if there is an additive or co-dependent role for *ERG3* and *FKS1* mutations in echinocandin resistance.

We performed controlled *in vitro* experiments with a drug-sensitive clinical isolate of *C. lusitaniae* to determine if MCF exposure alone is sufficient to select for multidrug resistance. We identified multiple mutations in genes involved in sterol biosynthesis, sterol organization, and cell wall biosynthesis under MCF exposure. We determined that MCF exposure alone is sufficient to select for single point mutants that have multidrug resistance, including *ERG3* and *OSH2* loss of function mutations. The multidrug resistance of the *ERG3* mutants is attributed to both changes in plasma membrane composition and cell wall remodeling through an absence of ergosterol, an increase in non-toxic alternative sterols, and increased chitin. This is the first conclusive evidence that mutations in *ERG3* are sufficient to cause multidrug resistance in *C. lusitaniae* and can evolve under echinocandin monotherapy.

## RESULTS

### Micafungin selection *in vitro* recapitulates within-patient evolution

To understand how MCF impacts the spectrum and dynamics of adaptive mutations in *C. lusitaniae*, we performed controlled *in vitro* evolution experiments in the presence and absence of MCF with the Day 1 pan-susceptible *C. lusitaniae* bloodstream isolate AMS5200 obtained from our previous case report (64). AMS5200, herein defined as the “progenitor” had the following minimum inhibitory concentrations (MIC): 0.016 µg/mL (MCF); 0.5 µg/mL (FLC); 0.094 µg/mL (AMB). We isolated six independent single colonies (designated A-F) from the progenitor, cultured them overnight in rich medium (YPAD), and subdivided each culture into three experimental conditions: 1) YPAD with no antifungal drug (“YPAD” lineages); 2) YPAD plus a constant, low concentration of MCF (0.016 µg/mL MCF; “Low” lineages); and 3) YPAD plus the same low concentration of MCF for passages 1-5, followed by an increase to 0.032 µg/mL MCF for passages 6-10 (“High” lineages). This increase in MCF concentration mirrors the increased dosage administered to the original patient halfway through MCF monotherapy (64). All lineages were diluted 1:1000 into fresh media every 24 hours for 10 passages, and cells were archived every passage for genotypic and phenotypic analyses. By passage 10 of the experiment, four MCF lineages went extinct including one Low MCF lineage (Lineage F) and three High MCF lineages (Lineages A, B, F) (Fig. 1A).

**FIG 1.**
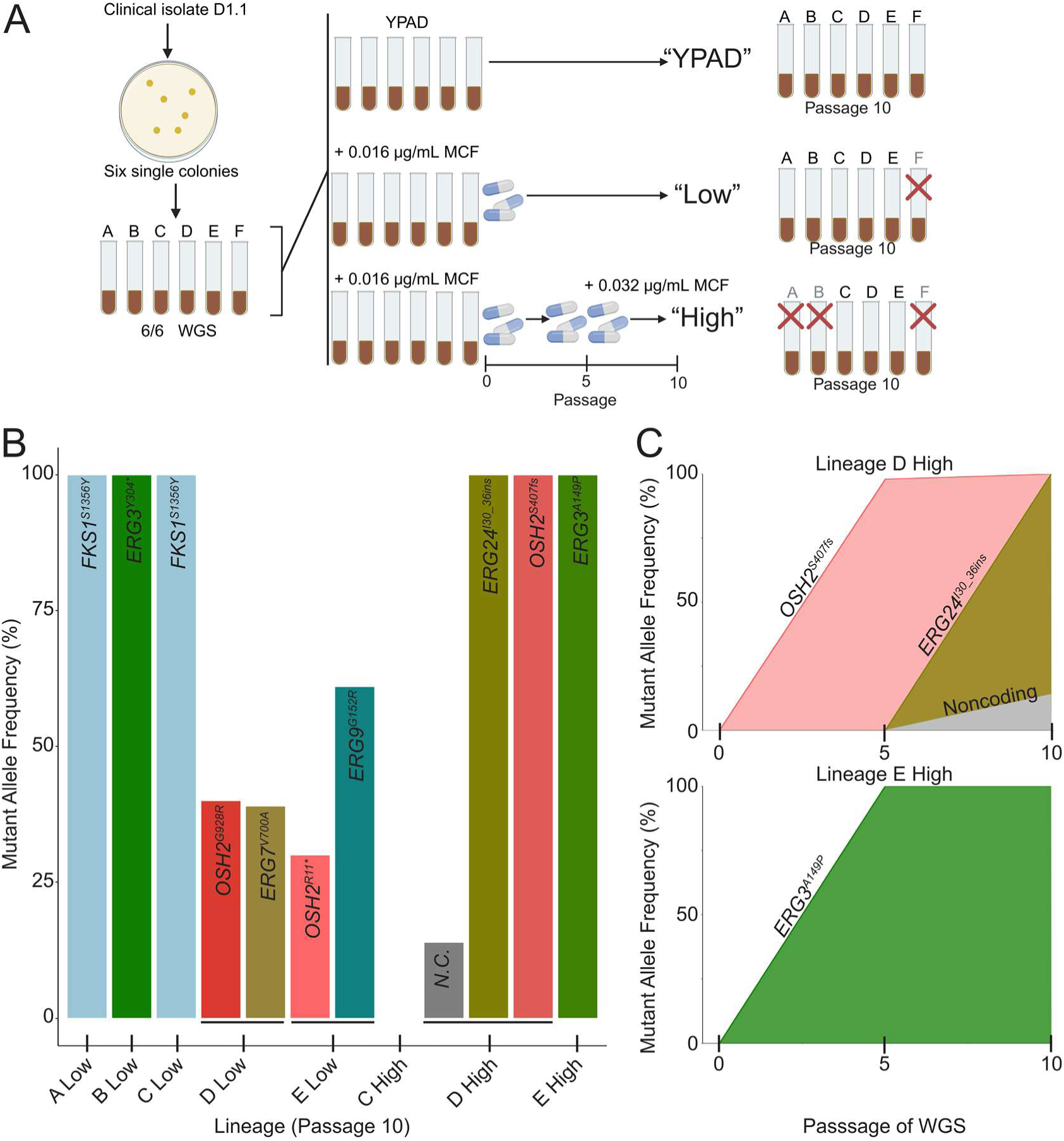
Mutations detected during micafungin exposure. (**A**) Schematic of the micafungin (MCF) evolution experiment. Six independent single-colony lineages (A-F) were derived from the drug-sensitive *C. lusitaniae* bloodstream isolate AMS5200 (64). All initial lineages were whole genome sequenced (WGS). Each lineage was passaged in three separate conditions: 1) YPAD for 10 passages; 2) YPAD + 0.016 µg/mL MCF for 10 passages (constant “Low” drug); and 3) YPAD + 0.016 µg/mL MCF for 5 passages followed by YPAD + 0.032 µg/mL MCF for another 5 passages (“High” drug). Populations that went extinct indicated with a red X. (**B**) Allele frequency of *de novo* sequence mutations identified from population-level whole genome sequencing from MCF evolved lineages at passage 10. Genes and one non-coding mutation (N.C.) that acquired mutations are indicated. (**C**) Mutant allele frequencies at passages 0, 5, and 10 for lineages D-High and E-High.

To identify genomic changes that arose during the evolution experiment, we performed whole genome sequencing on all six initial single colony populations (A-F) and all surviving populations at passage 10. Additionally, we generated a *de novo* long-read genome assembly of the progenitor to improve mapping quality and variant calling for the evolved populations (See Methods). We did not detect any DNA copy number changes in either the presence or absence of MCF (Fig. S1). In the absence of MCF, we detected no point mutations or small insertions/deletions relative to the progenitor. In contrast, the 5 surviving Low MCF lineages acquired 7 total mutations (1-2 mutations per lineage), whereas the 3 surviving High MCF lineages acquired 0-3 mutations per lineage, with no mutations detected in lineage C-High after additional variant calling and manual analysis (Fig. 1B, Table S2). Collectively, MCF selection resulted in *de novo* missense or nonsense mutations in *FKS1*, *ERG3*, *OSH2*, *ERG7*, *ERG9*, *ERG24*, and one non-coding region. Notably, several *FKS1* and *ERG3* mutations occurred within four amino acids of those identified in isolates obtained from the original patient receiving MCF monotherapy, with mutations repeatedly affecting the same domains (Fig. S2). For example, *FKS1^S1356Y^* and *FKS1^L1355S^*both occurred in the hotspot 2 domain associated with echinocandin resistance in *Candida* species (76, 77). Similarly, *ERG3^Y304*^* and *ERG3^Q308K^* occurred in the fatty acid hydroxylase domain of *ERG3*, which has been associated with multidrug resistance in clinical isolates of *C. lusitaniae* (64, 78). Collectively, our findings demonstrate that MCF exposure is sufficient to select for single nucleotide variants in *ERG3* or *FKS1*, both during *in vitro* selection and within a patient receiving MCF monotherapy.

### Resistance-associated mutations repeatedly reach fixation under micafungin selection

The recurrent but independent mutations in *FKS1* and *ERG3* indicate that these genes represent key adaptive nodes in response to MCF selection. *FKS1* and *ERG3* mutations occurred in four independent lineages and all reached fixation (100% allele frequency) with no additional mutations observed in the population (Fig. 1C, Fig. S3, Table S2). Identical *FKS1^S1356Y^* mutations arose in two independent Low MCF lineages (A-Low and C-Low), and different *ERG3* mutations, *ERG3^Y304*^* and *ERG3^A149P^*, arose in two independent MCF lineages (B-Low and E-High). Two other mutations reached fixation in lineage D-High, a frameshift in *OSH2^S407fs^* and a 7 amino acid in-frame duplication in *ERG24^I30_36ins^*, indicating that individual cells in this population harbor both mutations. Osh2 is a conserved oxysterol-binding protein that regulates intracellular sterol distribution via non-vesicular transport at membrane contact sites (79–81) and Erg24 is a C-14 sterol reductase late in ergosterol biosynthesis which is not essential in *C. albicans* (82–84).

Additional mutations occurred at intermediate (<100%) frequencies in *OSH2*, *ERG9*, *ERG7*, and a non-coding locus, and were detected exclusively in lineages with at least one additional mutation (lineages D-Low, E-Low, and D-High). Lineage D-Low acquired *ERG7^V700A^*and *OSH2^G928R^* mutations at 39% and 40% frequency. *ERG7* encodes the lanosterol synthase enzyme early in ergosterol biosynthesis (85). Similarly, lineage E-Low acquired *ERG9^G152R^*and *OSH2^R11*^* mutations at 61% and 30% frequency. *ERG9* encodes squalene synthase, the first committed enzyme in ergosterol biosynthesis (86). To determine if mutations that reached intermediate frequencies arose in distinct subpopulations, we isolated at least five single colonies from the populations where more than one mutation was identified at an intermediate frequency and performed Sanger sequencing across each mutation (Tables S1 & S2). All single colonies contained only a single mutation, indicating that mutations rose to intermediate frequencies in separate subpopulations.

To understand how the High MCF populations were impacted by the increase in MCF concentration at passage 5, we performed additional comparative genomics at this passage for the surviving lineages. At passage 5, both D-High and E-High acquired a single high-frequency mutation (*OSH2^S407fs^* at 98% and *ERG3^A149P^* at 100%) that remained fixed in the population at passage 10. In lineage D-High, only the *OSH2 ^S407f^*^s^ mutation was present at passage 5, indicating that the *ERG24^I30_36ins^*and non-coding mutations arose after the increase in MCF concentration (Fig. 1C, Fig. S3).

### Micafungin selects for point mutations that confer a fitness benefit across all three antifungal classes

To evaluate whether MCF-selected point mutations conferred a fitness advantage, we measured antifungal susceptibility across the three major antifungal drug classes (MCF, FLC, and AMB). We used standardized MIC assays (stationary cultures) to capture clinically relevant resistance and growth-curve assays (shaking cultures) to quantify growth in the *in vitro* environment from which the mutants were selected, particularly for oxygen-dependent growth phenotypes such as sterol biosynthesis. We isolated single colonies from the evolved populations and confirmed their genotypes with Sanger sequencing. Nine mutants were selected to represent the following genotypes from independent lineages: *FKS1^S1356Y^-1* (from A Low), *FKS1^S1356Y^-2* (from C Low), *ERG9^G152R^*, *ERG7^V700A^*, *ERG24^I30_36ins^/OSH2^S407fs^*(double mutant), *OSH2^G928R^*, *OSH2^R11*^*, *ERG3^A149P^*, *ERG3^Y304*^* (Table S2).

All MCF selected mutants had an increase in MIC in at least one drug, and the *ERG3* mutants had the highest increase in MIC to all three antifungal classes. In MCF, the *FKS1* and *ERG3* mutants had ∼32-fold higher MIC than the progenitor (Fig. 2A; Table S3). In FLC, the *ERG3* mutants had a 32- to 128-fold higher MIC than the progenitor (Fig. 2B). In AMB, all *OSH2* and ergosterol biosynthesis mutants had a ∼2- to 17-fold higher MIC relative to the progenitor (Fig. 2C). Overall, the MIC assay indicated that the MCF selected point mutants acquired cross-resistance to FLC and AMB, despite no prior exposure to these drug classes.

**FIG 2.**
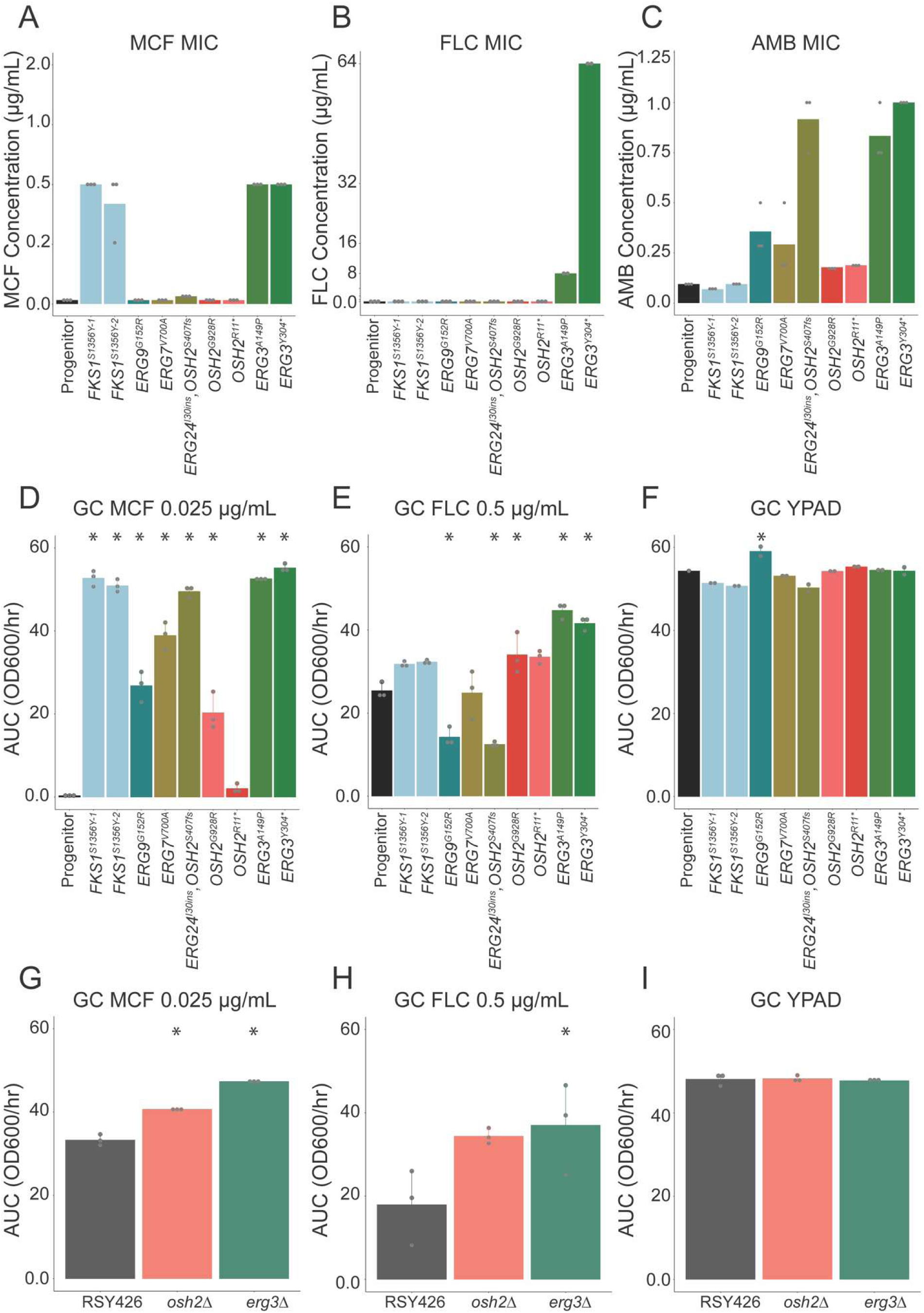
Micafungin-selected mutants have increased growth in all three antifungal classes, recapitulated by engineered deletions. Minimum inhibitory concentrations (MIC_50_) determined by the static broth microdilution assay for (**A**) MCF (0-8 µg/mL), (**B**) FLC (0-256 µg/mL), and (**C**) by E-test for AMB (0-32 µg/mL). All MIC values available in Table S3. Replicates are indicated by gray points. (**D-F**) 96-well growth curve analysis of the *in vitro* evolved mutants and progenitor. Area Under the Curve (AUC) measured by OD_600_ from growth curve assays in: (**D**) YPAD + 0.025 µg/mL MCF; (**E**) YPAD + 0.5 µg/mL FLC; or (**F**) YPAD. Replicates are indicated by gray points. Asterisks represent p < 0.05 relative to progenitor. Significant differences determined by ANOVA with Tukey HSD post-hoc analysis with adjustments for multiple comparisons (Table S3 for p-values). (**G-I**) Null mutants engineered in the RSY426 genetic background. AUC measured by OD_600_ from growth assays in: (**G**) YPAD + 0.025 µg/mL MCF (*osh2Δ*, *p = 0.0001; erg3Δ*, *p = 0.0*); (**H**) YPAD + 0.5 µg/mL FLC (*osh2Δ, p = 0.08; erg3Δ, p = 0.03*); or (**I**) YPAD. Asterisks represent p < 0.05 relative to RSY426. Significant differences determined by ANOVA with Tukey HSD post-hoc analysis and adjustments for multiple comparisons (Table S3).

The growth-curve assay revealed enhanced growth phenotypes for all mutants in the presence of antifungal. Apart from *OSH2^G928R^*, the mutants displayed significantly higher growth than the progenitor in 0.025 µg/mL MCF, a concentration intermediate between the Low and High selective environments (Fig. 2D). The higher growth in MCF supports that the mutations in *OSH2*, *ERG7*, *ERG9*, *ERG24*, and *ERG3* can confer a fitness advantage in the presence of drug, when cultures are shaking and well-oxygenated. In FLC, six of the nine mutants exhibited higher growth than the progenitor (Fig. 2E), with the *OSH2^G928R^* and both *ERG3* mutants showing statistically significant increases in growth (Table S4). In contrast, the *ERG24^I30_36ins^/OSH2^S407fs^* double mutant and the *ERG9^G152R^* mutant had reduced growth in FLC, indicating a context-dependent tradeoff associated with their adaptation to MCF. Notably, in the absence of drug, none of the mutants had reduced growth, and the *ERG9^G152R^* mutant had significantly increased growth relative to the progenitor (Fig. 2F). Overall, these data support that adaptation to MCF can select for independent point mutations that enhance fitness across distinct antifungal classes with different mechanisms of action without incurring a fitness cost.

### Erg3 or Osh2 loss-of-function alone causes multidrug resistance in *C. lusitaniae*

We repeatedly identified single point mutations in *ERG3* and *OSH2* in independent MCF selected lineages, suggesting these mutations confer a selective benefit in MCF. To determine if deletion of *ERG3* or *OSH2* is sufficient to cause multidrug resistance, we deleted each gene from the drug-sensitive *C. lusitaniae* strain RSY426 (*ku70::FRT, lig4::FRT*) that is deficient for non-homologous end joining (87). We confirmed the gene deletions by PCR and WGS (Table S1) and identified only a single off-target noncoding variant in the *erg3Δ* strain that arose during the transformation process. Deletion of *OSH2* or *ERG3* significantly increased growth in 0.025 µg/mL MCF compared to RSY426 (*osh2Δ, p =* 0.0001; *erg3Δ, p* < 0.0001) (Fig. 2G). In 0.5 ug/ml FLC, deletion of *ERG3* significantly increased growth (*p =* 0.03), while deletion of *OSH2* had a modest increase (*p* = 0.08, Fig. 2H). Neither deletion affected growth in the absence of drug (Fig. 2I). Together, these data indicate that loss of *OSH2* or *ERG3* confers a growth advantage under multiple antifungal classes.

The role of Erg3 loss of function in echinocandin resistance has been attributed to epistatic interactions with an Fks1 polymorphism in the *C. parapsilosis* species complex (71, 75, 88, 89). To determine if standing genetic variation within the echinocandin drug target *FKS1* could be contributing to the *ERG3* or *OSH2* resistance phenotypes in our *C. lusitaniae* lineages, we performed a multiple sequence alignment of the Fks1 protein sequence from our progenitor and five strains with distinct genetic backgrounds of *C. lusitaniae* (Fig. S4). Three of the five diverse genetic backgrounds had identical Fks1 sequences, and none had the P660A polymorphism that is fixed throughout the *C. parapsilosis* species complex (90), supporting that *de novo ERG3* mutations are sufficient to cause echinocandin resistance independent of an underlying *FKS1* polymorphism.

### Multidrug resistant *ERG3* mutants have major changes to cell wall and plasma membrane components

In *C. albicans,* deletion of individual ergosterol biosynthesis genes leads to hypersensitivity to Hsp90 inhibition (91). In *C. albicans* wild-type cells, Hsp90 inhibition reduces tolerance to MCF and FLC (91, 92). However, in *C. auris*, Hsp90 inhibition is not synergistic with FLC, suggesting a species-specific dependence on Hsp90 (93). To test whether resistance of *C. lusitaniae ERG3* point mutants requires Hsp90 activity, we quantified growth in the presence and absence of the Hsp90 inhibitor, radicicol. In addition to the two *in vitro* evolved *ERG3* mutants, we included the Day 9 patient isolate that evolved multidrug resistance during echinocandin monotherapy and is isogenic to the progenitor except for a single coding mutation in *ERG3^Q308K^* (64).

In YPAD and MCF, radicicol caused a significant reduction in growth for both *in vitro* evolved mutants (*ERG3^A149P^*, *p* = 0.006 (YPAD) and *p* = 0.009 (MCF); *ERG3^Y304^*, p* = 0.026 (YPAD) and *p* = 0.049 (MCF) (Fig. 3A&B, Table S4). In FLC, radicicol caused a 11-16% reduction in growth for all *ERG3* mutants. Radicicol did not significantly affect the growth of the progenitor in any condition (Fig. 3A-C, Table S4). In sum, *C. lusitaniae ERG3* mutants are more susceptible to Hsp90 inhibition in the absence of antifungal drugs than the wild-type progenitor, and depletion of Hsp90 improves the efficacy of both echinocandin and azole drugs against the *ERG3* mutants.

**FIG 3.**
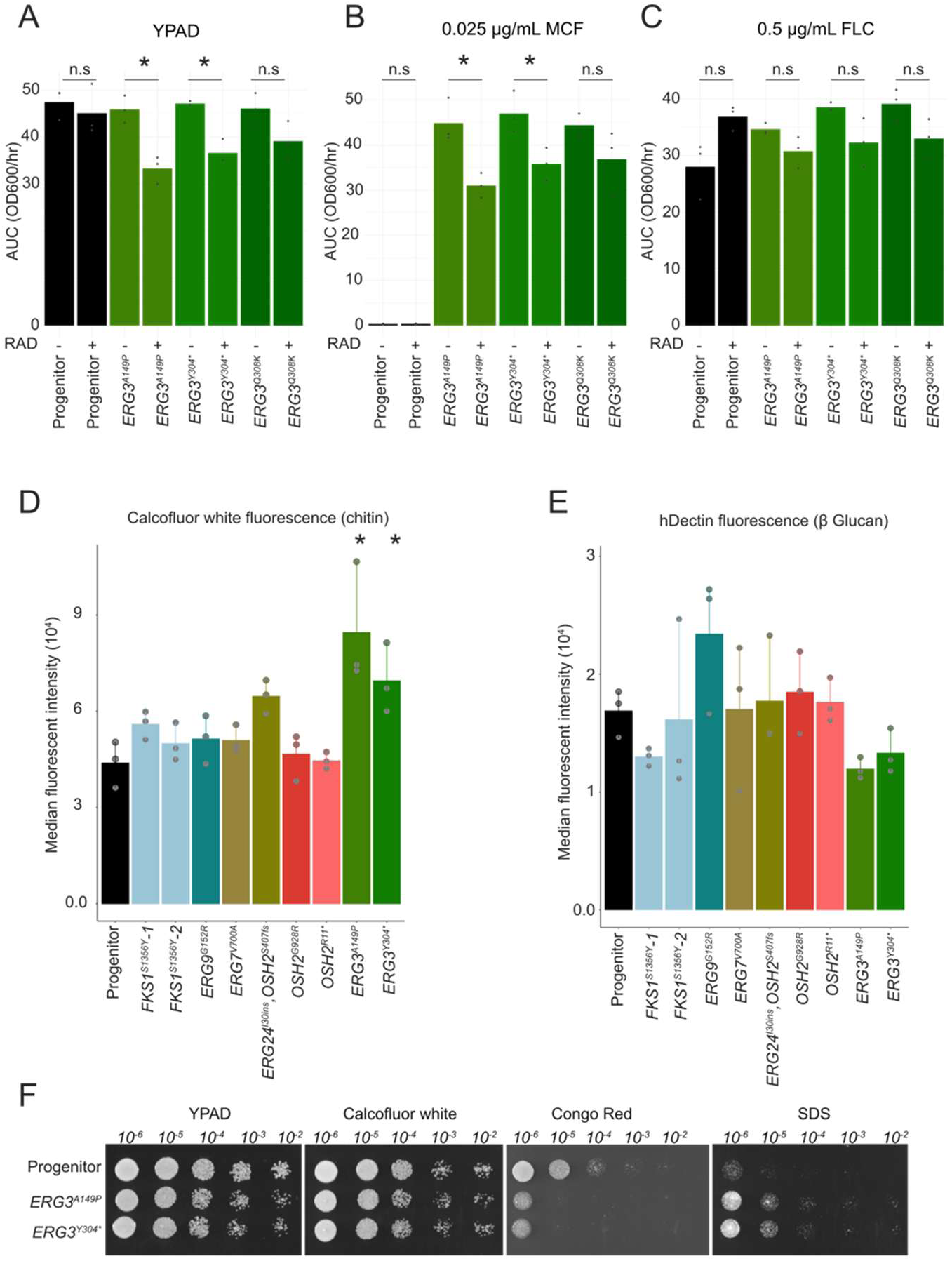
*ERG3* mutants are Hsp90-dependent with increased abundance of cell wall polymers. Area Under the Curve (AUC) measured by OD_600_ from 48 hour growth curve assays in the presence and absence of the Hsp90 inhibitor, radicicol (RAD) for the *C. lusitaniae* progenitor, evolved *ERG3* point mutants, and clinical isolate with *ERG3* mutation. (**A**) AUC in YPAD in the absence (-) or presence (+) of 2.5 µg/mL RAD. (**B**) AUC in 0.025 µg/mL MCF in the absence (-) or presence (+) of 2.5 µg/mL RAD. (**C**) AUC in 0.5 µg/mL FLC in the absence (-) or presence (+) of 2.5 µg/mL RAD. Gray points indicate technical replicates. Significance (p < 0.05, ANOVA with Tukey HSD) indicated with asterisks or ‘n.s.’ for not significant (p ≥ 0.05) (Table S4). (**D, E**) Median fluorescence intensity of (**D**) Calcofluor white that stains chitin (10 µg/mL) or (**E**) hDectin that stains 1,3-β-D-glucans (15 µg/mL) measured by flow cytometry (n ≥ 10,000 cells per isolate per replicate). Asterisk represents p < 0.05 relative to progenitor determined by ANOVA with Tukey HSD post-hoc analysis and adjustments for multiple comparisons (Table S5). (**F**) Serial dilutions (10^6^-10^2^) of evolved *ERG3* mutants and progenitor isolate were spotted on agar plates with YPAD or YPAD + 15 µg/mL Calcofluor white, + 150 µg/mL Congo Red, or + 0.005% Sodium Dodecyl Sulfate (SDS). All plates were repeated in triplicate, incubated at 30°C and imaged at 24 hours.

Given the role Hsp90 in cellular responses to cell wall stress and regulating sterol homeostasis, we tested whether the *ERG3* have alterations in cell surface molecules (94, 95). We quantified the abundance of major inner cell wall polymers by measuring fluorescence intensity of Calcofluor white (chitin) and hDectin antibody staining (1,3-β-D-glucan). The *ERG3* point mutants and the *erg3Δ* had significantly increased Calcofluor white fluorescence, indicating an increase in chitin abundance relative to the drug-sensitive progenitor (*ERG3^A149P^*, *p =* 0.0004, *ERG3^Y304*^*, *p =* 0.04, *erg3Δ*, *p =* 0.04, Fig. 3D, Fig. S5A). All *ERG3* mutants had decreased hDectin fluorescence (1,3-β-D-glucan) and increased sensitivity Congo Red (Fig. 3E&F, Fig. S5B&S6). Interestingly, *ERG3* mutants were less sensitive to the detergent SDS than the progenitor, indicating that the plasma membrane components might be altered (Fig. 3F) (96).

### *ERG3* mutants have a unique sterol profile consistent with Erg3 loss-of-function

Echinocandins do not directly target sterol biosynthesis and their impact on sterol composition in *Candida* species has not been investigated. We used GC-MS to determine the sterol composition of two *ERG3* point mutants (*ERG3^A149P^* and *ERG3^Y304*^*) relative to the progenitor in three separate growth conditions: no drug, 0.016 µg/mL MCF, and 0.5 µg/mL FLC.

First, using a targeted GC-MS approach we quantified the abundance of known sterols with commercially available standards (Fig. S7, Table S6). In the absence of antifungals, both *ERG3* mutants had significantly less ergosterol than the progenitor (*ERG3^A149P^*, *p* = 0.018; *ERG3^Y304*^*, *p* = 0.020). Ergosterol comprised 71% of the relative sterol abundance in the progenitor, but only <1% in the *ERG3* mutants, consistent with a functional loss of Erg3 (Fig. 4A, Table S6). The absence of ergosterol was also consistent with the AMB resistance identified in the *ERG3* mutants (Fig. 2C). In the presence of FLC, the progenitor had a significant increase in the abundance of ergosterol intermediates obtusifoliol and lanosterol (*p* = 0.00096), associated with increased flux through the alternative pathway (Fig. 4A, Fig. 4D). FLC exposure in the *ERG3*^A149P^ mutant led to a significant increase in zymosterol (*p* = 0.0007) and significant decrease in episterol (*p* = 0.00005), the precursor to the *ERG3*-specific sterols ergosta-7,22-dienol and ergosta-7-enol. However, FLC exposure in the *ERG3* mutants did not suggest increased flux through the alternative pathway, with no significant accumulation of the alternative pathway intermediates (Fig. 4D, Table S6). In MCF, the progenitor had a decrease in ergosterol and an increase in the alternative sterol species (Fig. 4B). MCF affected the two *ERG3* mutants differently: the *ERG3^A149P^* mutant had only a significant increase in zymosterol in the presence of MCF (*p = 0.016*, Table S6), whereas the *ERG3^Y304*^*mutant had increased alternative and canonical pathway sterols, including a significant increase in eburicol, in the presence of MCF (*p* = 0.0005; Fig. 4D, Table S6).

**FIG 4.**
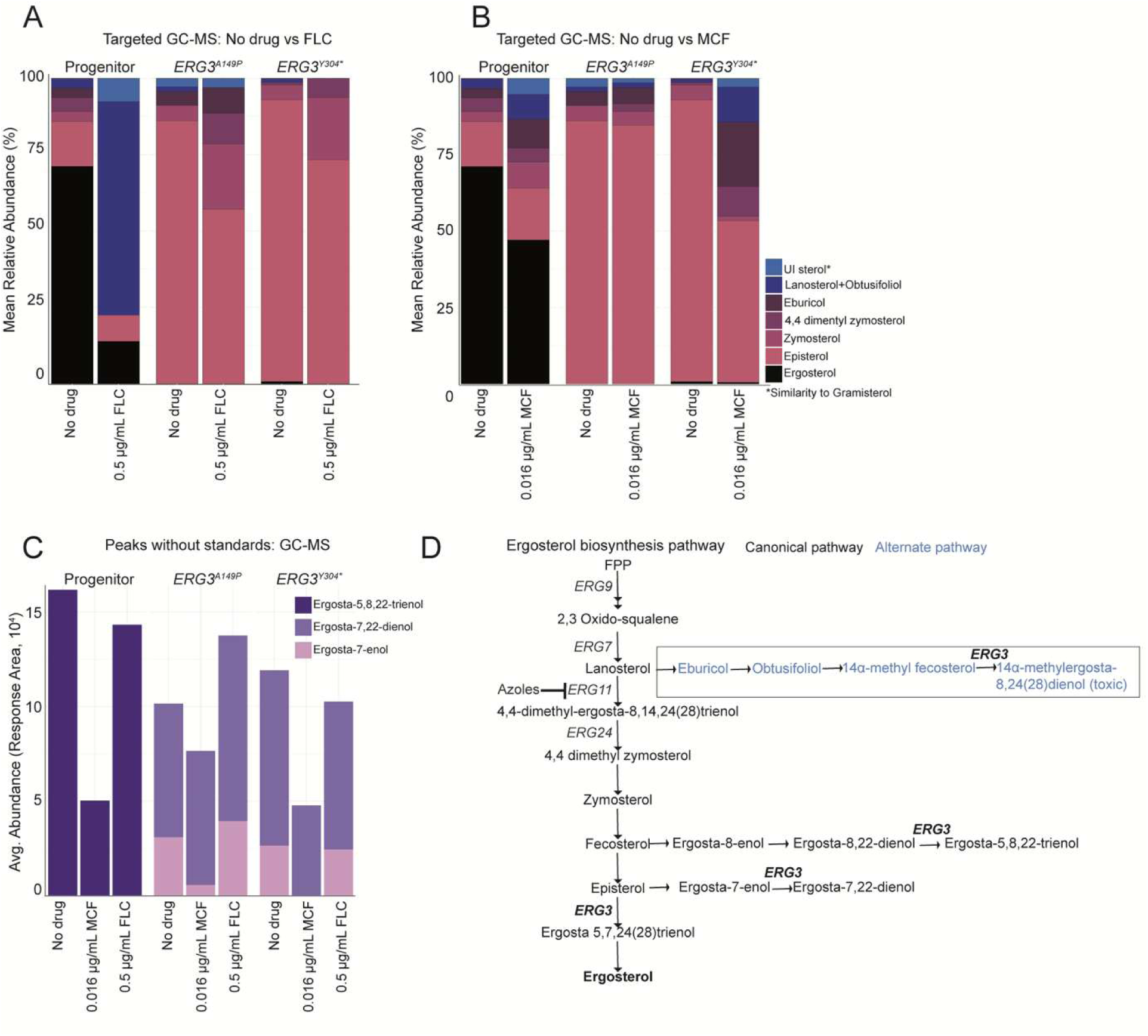
*ERG3* mutants have a unique sterol profile consistent with Erg3 loss of function. (**A**) Mean relative abundance of sterols identified using targeted GC-MS after growth in the absence of antifungals or after growth + 0.5 µg/mL FLC. Mean value was calculated using three biological replicates per strain and per condition. See Table S6 for p-values. Asterisks indicate unidentified sterol that has similarity to Gramisterol. (**B**) Mean relative abundance of sterols identified using targeted GC-MS after growth in the absence of antifungals or after growth in 0.016 µg/mL MCF. Mean value was calculated using three biological replicates per strain and per condition. See Table S6 for p-values indicating significant differences in abundance of sterols. (**C**) Average response area of sterols without standards identified using GC-MS analysis after growth in the absence of antifungals or after growth + 0.016 µg/mL MCF or 0.5 µg/mL FLC. (**D**) Ergosterol biosynthesis pathway inferred from (48,98,68,97). Blue font indicates sterols synthesized as part of the alternative pathway, initiated following azole inhibition of Erg11. Biosynthetic steps where Erg3 functions are indicated in bold.

Second, on observing the chromatograms we identified two unknown sterol species that were exclusive to the *ERG3* mutants (Fig. 4C), with highest matches to the episterol derivatives ergosta-7,22-dienol and ergosta-7-enol, that have been reported in *ERG3* mutants of *C. albicans* (Fig. 4D, Fig. S8) (67, 68, 97). Similarly, we predicted one unknown sterol exclusive to the progenitor in all three conditions to be ergosta-5,8,22-trienol, a fecosterol derivative that is synthesized by Erg3 from an ergosta-8,22-dienol precursor (Fig. 4D, Fig. S8). In summary, FLC and MCF exposure was sufficient to cause significant changes the sterol profiles of *C. lusitaniae* wild-type cells and these changes were dominated by a loss of ergosterol and increase in alternative sterols. In contrast, the sterol profiles of the *ERG3* mutants were dominated by episterol and its derivatives, in both the absence and presence of FLC and MCF.

## DISCUSSION

Despite increasing reports of multidrug resistance in non-*albicans Candida* species, the molecular mechanisms and evolutionary trajectories driving multidrug resistance remain poorly characterized (99, 100). We found that exposure to a single echinocandin drug is sufficient to select for resistance to all three classes of antifungals in *C. lusitaniae*. The echinocandin MCF recurrently selected for mutants with a single, *de novo* point mutation in *ERG3*, *FKS1*, or *OSH2*. We provide evidence that *erg3* and *osh2* loss-of-function mutations are sufficient to confer fitness benefits in multiple classes of unrelated antifungals independently of genetic background. The current *in vitro* study parallels our previous case report describing the evolution of a *de novo* multidrug resistant *ERG3^Q308K^* mutation in a patient receiving echinocandin monotherapy over an 11-day period with no prior history of azole therapy (64). Three independent *FKS1* mutations also evolved in this patient, but all were independent of the *ERG3* mutant genotype. Similarly, two studies of other *Candida* species report individual *ERG3* mutations that caused azole and echinocandin cross-resistance, including *ERG3^G111R^* in a *C. parapsilosis* clinical isolate (71), and *ERG3^Y325*^* and *ERG3^Y190*^* nonsense mutations in *C. albicans* clinical isolates (101), underscoring that *ERG3* mutations are relevant to mechanisms of in multidrug resistance across diverse fungal species.

The independent role of *ERG3* mutations in echinocandin resistance has been debated because *ERG3* polymorphisms frequently occur in genetic backgrounds with other mutations, including *FKS1* polymorphisms, across diverse *Candida* species (71, 75). For example, the *C. lusitaniae ERG3^Q308*^* mutation was identified in a candidemia patient treated with a complex antifungal regime that included the echinocandin caspofungin, however, the link between *ERG3^Q308*^* and multidrug resistance was not considered because several additional mutations, including *FKS1* and *ERG4*, were present in this isolate (78). Additionally, the contribution of *ERG3* mutations to antifungal resistance has species-specific differences. For example, there is a fixed *FKS1* polymorphism in the *C. parapsilosis* species complex that likely reduces echinocandin susceptibility in general (75, 90, 102). Importantly, we found no evidence for concurrent *FKS1* polymorphisms in any of our *de novo ERG3* mutants nor fixed variants across five diverse genetic backgrounds of *C. lusitaniae*, and the *de novo ERG3* mutations all arose in drug sensitive backgrounds. The Erg3 enzyme from different fungal pathogens, including *C. albicans*, *N. glabratus*, *C.auris*, *Cryptococcus neoformans*, *Aspergillus fumigatus*, and *Rhizopus delemar* also confer variable susceptibilities to fluconazole, indicating a conserved but not identical role for this protein across species (19). Notably, *ERG3* deletion alone in *C. albicans*, *C. auris*, and *N. glabratus* does not cause echinocandin resistance, and *ERG3* point mutations can even increase sensitivity to echinocandins when combined with *erg11* deletion (75, 103–105). We found that deletion of *ERG3* in a drug-sensitive *C. lusitaniae* background recapitulated the multidrug resistant phenotypes of the *ERG3* mutants obtained from the patient and *in vitro*, confirming that loss of *ERG3* is sufficient to confer multidrug resistance in this species. More research is needed to determine the mechanisms driving the species-specific phenotypes of Erg3, including a deeper understanding of the functional role of different polymorphisms in the presence and absence of antifungal drugs.

We found that multidrug resistance can emerge under single-drug selection with the echinocandin MCF in *C. lusitaniae*. In *N. glabratus*, similar *in vitro* exposure to the echinocandin anidulafungin selected for *ERG3* mutations associated with echinocandin and azole cross-resistance, however these mutations occurred concurrently with mutations in *FKS1*, *FKS2*, or *CNE1* and causal mechanisms were not verified (106, 107). Interestingly, exposure to an azole drug did not result in the same cross-resistance in *N. glabratus* (107), indicating that echinocandin exposure is both necessary and sufficient to select *ERG3*-linked multidrug resistance. Given the widespread clinical use of echinocandins, these findings highlight a critical risk: treatment with a single echinocandin might inadvertently select for broad antifungal resistance, thereby constraining the already limited therapeutic options.

Echinocandin drugs are lipopeptides that target Fks1 directly in the lipid membrane (108, 109). Cryo-electron microscopy and deep mutational scanning indicate that Fks1 polymorphisms can inhibit drug binding and confer resistance, and that these drug interactions are occurring directly in the membrane (31, 71, 110, 111). Because of this, changes in sterol composition are proposed to impact the interaction between Fks1 and echinocandin drugs. Strikingly, we found that echinocandin exposure was sufficient to affect the sterol composition of wild-type *C. lusitaniae* cells. The altered sterol composition in the presence of MCF could then be selected on. While we identified two independent lineages that acquired *FKS1* mutations, the majority of the mutations identified under MCF selection were within ergosterol biosynthesis genes (*ERG3*, *ERG24*, *ERG9*, *ERG7*) or sterol transporters (*OSH2*). Ergosterol depletion and increased abundance of alternative sterols like zymosterol mediate MCF resistance. Zymosterol accumulates in the *ERG3^A149P^*mutant under MCF exposure and similarly accumulates in a *C. albicans osh2Δ/Δ* mutant (75), indicating that the balance and composition of membrane sterols are important for conferring antifungal resistance. Interestingly, no mutations were identified in one MCF selected lineage (Lineage C High), despite rigorous comparisons for all types of sequence variants. One possible explanation as to how this lineage survived without acquiring point mutations could be through epigenetic modifications. In *C. auris*, loss of histone H3 acetylation through depletion of the acetyltransferase *GCN5* downregulates ergosterol biosynthesis genes and affects cell wall integrity, highlighting that non-genetic mechanisms can mediate mechanisms that confer antifungal resistance (112). More studies are needed to understand the diversity of genetic and non-genetic mechanisms that can promote adaptation to echinocandin drugs.

Our model for multidrug resistance in *ERG3* mutants is dependent on the altered sterol profile. We show that the *in vitro* evolved *ERG3* mutants have phenotypes that indicate loss of function, including an absence of ergosterol and accumulation of alternative sterol species (Fig. 4A, 4C). Therefore, we posit that resistance to azoles and polyenes is consistent with previously reported models of *ERG3* loss of function: 14α-methylergosta-8,24(28)-dien-3,6-diol is not synthesized, conferring azole resistance, and ergosterol is not synthesized, conferring polyene resistance (15, 97). We build on this model by suggesting that the lipid profile also either directly or indirectly mediates echinocandin resistance. Alternative sterols exclusive to the *ERG3* mutants are sufficient to maintain phase separation in the absence of ergosterol (113). However, MCF incorporation into the plasma membrane is ergosterol-specific, where MCF cannot be incorporated into a membrane that contains cholesterol instead of ergosterol (114). While it is untested whether MCF can penetrate a membrane that contains ergosta-7,22-dienol and ergosta-7-enol instead of ergosterol, it could be the case that these sterols are sufficient for steric exclusion of MCF into the membrane, thus conferring some level of resistance. The replacement of ergosterol with ergosta-7,22-dienol and ergosta-7-enol might also indirectly induce signaling cascades that upregulate cell wall biosynthesis. Additionally, the absence of ergosterol can incur changes to chitin regulation (115–117). Increased chitin in *C. albicans*, *C. tropicalis*, *C. krusei* and *C. parapsilosis* was identified in isolates treated with fluconazole and concurrent with ergosterol depletion (118). The *HOG* pathway that regulates expression of chitin synthase genes and chitin synthesis in *C. albicans* is also activated under ergosterol depletion in *S. cerevisiae*, indicating a potential link for the regulation of cell surface composition (119, 120). We propose that even if echinocandins can permeate the *ERG3* mutant plasma membrane, the increased chitin abundance in the *ERG3* mutants would be sufficient to maintain cell wall integrity under Fks1 inhibition (121).

## CONCLUSION

Multidrug resistance evolves during MCF treatment both *in vitro* and during antifungal therapy in a patient. Mutations in sterol biosynthesis and sterol transport, primarily *ERG3* and *OSH2* loss of function mutations, conferred multidrug resistance and competed with canonical *FKS1* mutations in the presence of an echinocandin drug. Major changes in sterol and cell wall composition underlie the acquisition of antifungal resistance for *ERG3* mutations. The rapid emergence of multidrug resistance following exposure to a single antifungal agent highlights the urgency to test more frequently for the acquisition of resistance during antifungal treatment. Furthermore, it is necessary to consider the diversity of genetic drivers and mechanisms of resistance that could contribute to treatment failures, especially in rare and emerging pathogens.

## MATERIALS AND METHODS

### Strains used in this study

All whole genome sequenced strains and single colonies used in this study are listed in Table S1. Strains were stored at −80°C in 20% glycerol. All strains were cultured on YPAD medium (yeast extract, peptone, and 2% dextrose supplemented with 40 μg/ml adenine and 80 μg/ml uridine) and incubated for 48 hours at 30°C, unless otherwise noted.

### *In vitro* evolution

The drug-sensitive clinical isolate AMS5200 (“progenitor”) was cultured from frozen −80°C glycerol stock and struck onto YPAD agar plates for single colonies (64). Six single colonies from the progenitor were selected at random and suspended in 3 mL YPAD liquid and incubated ∼16 hours at 30°C. After overnight growth, each liquid culture was diluted 1:1000 into 3 mL of media in each of three treatment conditions: 1) YPAD only; 2) YPAD + 0.016 μg/ml MCF (constant “Low”); or 3) YPAD + 0.016 μg/ml MCF that then increased to 0.032 µg/mL MCF on passage 5 and continued for the reminder of the experiment (“High”) and incubated with constant agitation for 24 hours at 30°C. Every 24 hours (one passage), cells were vortexed to resuspend and diluted 1:1000 in 3 mL of fresh medium. In total, 10 passages were conducted, and at each passage, lineages were collected for storage in glycerol at −80°C.

### Long-read sequencing and *de novo* genome assembly of *C. lusitaniae* clinical isolate AMS5200

Library preparation for clinical isolate AMS5200 was performed using the PacBio SMRTbell® prep kit 3.0 with the SMRTbell® barcoded adapter plate 3.0. Sequencing was performed on a PacBio Sequel IIe for a 15-hour collection period, followed by demultiplexing, QC and adapter trimming. Reads were checked for adapter contamination with HiFiAdapterFilt (v2.0.1) and summary statistics were generated with NanoPlot (v1.41.6) (122, 123). Genome assembly was performed with Flye (v2.9.1-b1780, default parameters, estimated genome size 13m) (124). Cleaning, sorting, repeat masking and annotation was performed with Funannotate (v1.8.9), run interactively with Interproscan (v5.23-62.0), eggnog-mapper (v2.1.6) and phobius (v1.01) (125–128). Gene content was assessed with BUSCO (“Benchmarking Universal Single-Copy Orthologue”, v5.3.2, dataset saccharomycetes_odb10) (129). Assembly data can be accessed at NCBI Bioproject PRJNA231221 (https://www.ncbi.nlm.nih.gov/bioproject/231221).

### Whole genome sequencing and variant detection

Population-level Illumina whole genome sequencing was performed on lineages collected from original single colonies, passage 5, and passage 10 of the experiment. Genomic DNA was isolated using a phenol-chloroform extraction as previously described (130, 131). Sample libraries were prepared using the Illumina DNA Prep kit and IDT 10bp UDI indices, and sequenced on an Illumina NextSeq 2000, producing 2×151bp reads. Demultiplexing, quality control and adapter trimming was performed with bcl-convert (v3.9.3). Quality trimming was performed for all isolates using BBDuk (part of BBTools, v38.94) (132). Reads were aligned to the AMS5200 progenitor reference genome assembly using BWA-MEM (v0.7.17) (133). Aligned reads were sorted by coordinate, duplicates were marked and bam files indexed by samtools (v1.10) (134). Raw read and alignment quality were assessed with FastQC (v0.11.9), qualimap (v.2.2.2-dev) and samtools flagstat, and summarized with MultiQC (v1.16) (135–137). *De novo* SNPs and small indels were called with Mutect2 (GATK v4.1.2) (138). To identify variants in the initial colonies, the progenitor was labeled as normal, and the initial isolate was labeled as tumor (variant calling was performed independently for lineages A through F). To identify variants in each of the evolved lineages, the initial isolate was labeled as normal, and the related evolved lineage was labeled as tumor (e.g., initial isolate A was normal and lineage A “Low MCF” was labeled as tumor), repeating the process for each lineage and each evolved population for each treatment. To identify variants in each sequenced transformant, the RSY426 background strain was labeled as normal and the transformant was labeled as tumor. All variants that passed Mutect2 quality filters were then manually filtered with the following parameters: minimum 5 unique alt reads, allele frequency ≥ 0.1, not supplementary or secondary alignment, and supporting reads in both orientations. All variants passing manual filtering parameters were visually verified in IGV (139). Lineage C “High MCF” had no variants detected after manual filtering. Therefore, as a quality check, we searched for any variants in the lineage C initial isolate and lineage C “High MCF” that passed Mutect2 quality filters prior to manual filtering steps, but none were identified.

### Relative copy number analysis

Yeast Mapping Analysis Pipeline (YMAP) was used to visualize genome-wide copy number for all sequenced isolates relative to the *de novo* genome assembly of AMS5200 (140). Relative copy number was then determined using sequencing read depth of GC-corrected alignment files (141, 142). Samtools (v1.10) depth with the option -aa was used to compute the read depth of all positions, and the rolling mean was then calculated over 500 bp tiled windows using the R package *RccpRoll* (v0.3.0) (143, 144). The median nuclear genome depth was calculated, and the relative read depth was calculated by dividing the window’s read depth by the median depth.

### Domain homology in *C. lusitaniae* mutants

Translation of the annotated sequences from the progenitor assembly was completed using Benchling. Amino acid sequence of the translated *ERG3* gene body from the *C. lusitaniae* AMS5200 strain (chromosome 6, 250562-251839, BioProject PRJNA954073) and sequence of the translated *FKS1* gene body from the *C. lusitaniae* AMS5200 strain (chromosome 3, 1254106-259766, BioProject PRJNA954073) were aligned to the translation of the *C. albicans* SC5314 (A22 reference) and *C. auris* B8441 strains (145). A pairwise identity cutoff of 90% was used for domain identity. Based on this alignment, predicted domains from UniProt (https://www.uniprot.org/uniprotkb/O93875/entry#family_and_domains) were assigned to the *ERG3* and *FKS1* sequences.

### PCR and Sanger sequencing

To identify cells that harbored a low frequency mutation within the population, single colonies were isolated from frozen glycerol stocks onto YPAD plates. Plates were incubated at 30°C overnight, single colonies were randomly selected and transferred to 3 mL YPAD and incubated at 30°C overnight with shaking. Genomic DNA was extracted from the single colonies using a standard phenol-chloroform extraction protocol. Primers used to amplify the target genes were designed using NCBI PrimerBlast (Table S1). Genes of interest were amplified using PCR and verified using agarose gel electrophoresis (Table S1). PCR products were diluted and sent for Sanger sequencing (Azenta/GeneWiz). Presence of the mutation of interest was determined by manual comparison of the Sanger sequencing data with the *de novo* genome assembly of the AMS5200 progenitor using SnapGene. Single colony isolates with the verified mutations were stocked in 20% glycerol and stored at −80°C.

### MIC drug susceptibility testing

Minimum inhibitory concentration (MIC) values for both FLC (ThermoFisher; 86386-73-4) and MCF (MedChemExpress; 208538-73-2) were determined using the broth microdilution method (141). Overnight cultures were inoculated from frozen glycerol stocks into YPAD with 1% dextrose at 30°C and diluted to a final OD600 of 0.01. Aliquots of 20 µL of each diluted culture was inoculated into 96-well plates containing increasing concentrations of either drug, as well as a no-drug control of fresh media. In MCF, the concentrations of the drug ranged from 0 to 8 µg/mL in 2-fold serial dilutions. In FLC, the concentrations of the drug ranged from 0 to 256 µg/mL in 2-fold serial dilutions. Inoculated plates were stationary, incubated at 30°C in a humidified chamber. Cultures were resuspended and OD_600_ readings were measured at 24-h and 48-h post inoculation using a BioTek Epoch2 plate reader. The MIC_50_ values were reported as the lowest concentration of antifungal drug that inhibits ≥50% of growth at 24 hours relative to the no-drug control. A mean MIC value was calculated using three triplicate assays in each condition per sample.

### E-test method for Amphotericin B

Antifungal susceptibility to AMB E-test strips was adapted from the CLSI Performance Standards for Antifungal Susceptibility testing of Yeasts using gradient diffusion strips (146). Briefly, frozen stocks were struck out onto Sabouraud’s Dextrose Agar plates and incubated at 35°C for 24 hours. Cells were collected by scraping the plate and resuspended into 200 μl of PBS, then diluted to a normalized OD_600_ of 0.1 on a BioTek Epoch2 plate reader. 100 µL of the diluted inoculum were plated onto RPMI plates and spread with glass beads. Each strain was plated on a control plate with no E-strip as well as a plate with an AMB E-strip (Liolfilchem). The RPMI plates were incubated at 35°C and images were taken at 24 hours and 48 hours using the BioRad GelDoc. MIC values were assessed blindly by at least three different individuals and repeated for three biological replicates. MIC values were designated where the growth intersects with the testing strip, and the highest value was taken (146,147). The CDC strain AR398 was used as a control.

### Growth curve analysis

Overnight cultures were grown from frozen glycerol stocks in 3 mL YPAD at 30°C and diluted to a final OD_600_ of 0.01. 20 µL were added to a 96-well plate containing YPAD with 1% dextrose and one of the following: 0.5 µg/mL FLC, 0.016 µg/mL MCF, or a no-drug control. Plates were incubated at 30°C with constant shaking for 48 hours, with OD_600_ readings every 15 minutes using the BioTek Epoch2 plate reader. Growth curve assays were performed in triplicate for each condition. Radicicol (RAD) growth curves were performed as above with the addition of 0 µg/mL, 0.625 µg/mL, 1.25 µg/mL, or 2.5 µg/mL RAD. Data were analyzed using GrowthCurveR and visualized using the R package ggplot2 (148, 149). All average AUC values were calculated from three technical replicates and are reported in Table S4.

### Spot plates

Overnight cultures were grown from frozen glycerol stocks in 3 mL YPAD at 30°C with shaking and diluted to an OD_600_ of 1.0 (3 x 10^7^ cells/mL). Serial dilutions were performed by adding 20 µL of culture into 180 µL of PBS, and 5 µL (3 µL for SDS) of each dilution were spotted onto the agar plates. All agar plates were YPAD with or without a chemical stressor: 0.005% SDS (sodium dodecyl sulfate); 15 µg/mL Calcofluor white; and 150 µg/mL Congo Red. All plates were incubated at 30°C and images were taken at 24 and 48 hours using BioRad Gel Doc and ImageLab.

### Flow cytometry analysis of cell wall components

Methods were adapted from (150). Briefly, all strains struck out from frozen glycerol stocks onto YPAD agar plates and incubated overnight at 30°C. Single colonies were inoculated into 3 mL YPAD media and grown overnight at 30°C with constant agitation. After ∼16 hours of growth, cells were washed in 1 mL PBS three times. Samples were then vortexed on top speed for 30 seconds to break up any clumping. Unstained control strains were set aside. To stain for exposed 1,3-β-D-glucan, cells were first blocked with 3% bovine serum albumin and 5% normal goat serum (Invitrogen) for 30 minutes. After blocking, cells were stained with 15 µg/mL of 611 hDectin-1a (InvivoGen) for 1 hour in the dark. Cells were washed twice with PBS before secondary staining with 4 mg/mL of goat raised anti-human IgG antibody conjugated with Alexa Fluor 647 (Invitrogen) for 30 minutes in the dark. To quantify chitin content, cells were stained with 0.1 µg/mL of Calcofluor white (Sigma) for 10 minutes in the dark. After the Calcofluor white staining, cells were washed with 50 µL PBS three times and resuspended in 500 µL PBS. Samples were analyzed on the Cytek Aurora flow cytometer with a minimum of 10,000 singlets recorded for each sample. Fluorescent channels used were R3-A (hDectin) and V3-A (Calcofluor white). Data were analysed using FlowJo (https://www.flowjo.com/solutions/flowjo/downloads) (v.10.10.1)

### Clavispora lusitaniae transformations

*C. lusitaniae* transformations were adapted from a *C. auris* protocols (151) with minimal modifications. Transformations were performed using the *C. lusitaniae* RSY426 strain background (*ku70::FRT*, *lig4::FRT*) for increased transformation efficiency (87). The RSY426 strain was struck out from frozen onto YPAD plates and incubated at 30°C overnight. A single colony was isolated and inoculated into 5 mL of YPAD media and incubated at 30°C overnight with agitation. The following morning, the overnight culture was pelleted using centrifugation and resuspended into 8 mL H_2_O, 1 mL TE, and 1 mL Lithium Acetate (LiOAc), then incubated at 30°C for one hour. 500 µL of DTT was added and incubation continued for 30 min. Cells were pelleted by centrifugation, washed in cold H_2_O then resuspended in 200 µL of sorbitol. 45 µL of cells were added to pre-chilled 2 mm gap electrocuvette with the transformation cassette and kept on ice. Cells were pulsed using the fungi setting (1.5 kV, 1 pulse) on the MicroPulser Electroporator (BioRad), then rescued in 1 mL of sorbitol. Cells were pelleted and resuspended in 10 mL YPAD and incubated for 2 hours at 30°C with gentle agitation. Cells were then plated onto selective media and incubated at 30°C for 24-48 hours.

All plasmids used for transformation can be found in Table S1. All primers used for transformations can be found in Table S1. To engineer the *osh2*Δ and *erg3Δ* strains in the RSY426 background, linear transformation repair cassettes were amplified from pAS3108 using primers 1967 and 1970, and pAS3109 using primers 1961 and 1964. Transformations were performed as described above. Gene deletion was confirmed by PCR bridging across the deletion cassette from each flanking region. PCR validated transformants were then screened by whole genome sequencing, and variant calling was performed as described above.

Plasmid pAS3108 was created using the pUC19 cloning vector, a NAT (Nourseothricin) resistance cassette, and an upstream and a downstream homology region of the *OSH2* gene from the progenitor (AMS5200). Fragments were amplified using primers 1578 and 1579, 1577 and 1575, 1967 and 1968, and 1969 and 1970, respectively. The plasmid was assembled using the NEBuilder HiFi Assembly Kit (NEBiolabs) according to manufacturer instructions and sequence verified by whole-plasmid sequencing (Plasmidsaurus).

Plasmid pAS3109 was created using the pUC19 cloning vector, a NAT (Nourseothricin) resistance cassette, and an upstream and a downstream homology region of the *ERG3* gene from the progenitor (AMS5200). Fragments were amplified using primers 1578 and 1579, 1577 and 1575, 1961 and 1962, and 1963 and 1964, respectively. The plasmid was assembled using the NEBuilder HiFi Assembly Kit (NEBiolabs) according to manufacturer instructions and sequence verified by whole-plasmid sequencing (Plasmidsaurus).

### Sterol analysis

Strains were struck on YPAD agar plates from frozen stocks and incubated at 30°C for 24 hours. Cultures were inoculated in 50 mL YPAD and incubated at 30°C, 220 rpm for 16 hours. Overnight cultures were pelleted by centrifugation and then washed once with PBS. Cells were resuspended in 25 mL of PBS, then 500 µL of the resuspension was added to 50 mL of fresh, filter-sterilized CSM medium (1.7 g/L YNP (DIFCO), 5.0 g/L Ammonium Sulfate (Sigma), 20 g/L Glucose (Sigma), 2 g/L SC AA Mix (MP Biomedical)) containing 0.5 μg/mL FLC, 0.016 µg/mL MCF, or no drug, and incubated at 30°C for another 5–6 hours with shaking. Cell density was quantified and 5×10^8^ cells were harvested by centrifugation, resuspended in 1 mL of PBS, then centrifuged again with removal of supernatant, and the pellet was frozen in −80°C. Lipid extraction was conducted as previously described (152). Briefly, pellets with 5×10^8^ cells were subjected to Mandala extraction and the Bligh & Dyer method for extraction of lipids. 1/4^th^ of the sample was aliquoted for the determination of inorganic phosphate; the remaining sample was base hydrolyzed and dried. The dried total samples were resuspended in 100 μL of BSTFA reagent (Thermo Scientific) and incubated at 70°C for 1 hour. Cholesterol (Avanti catalog # 700100) was added as an internal standard for these analyses prior to lipid extraction. The ramp, temperature, and hold time can be found in Table S6.

GC–MS sample preparation, analysis, and data processing: All derivatized samples were stored at −80°C upon receipt until analysis at the Stony Brook Proteomics Core Facility. Samples were analyzed on an Agilent 7890B GC coupled to a 5977A MS system and equipped with an autosampler. A 1 µL aliquot was injected in split mode (2:1) at 280°C. High-purity helium served as the carrier gas at a constant 1.2 mL/minute flow. Separation was performed on an HP-5MS capillary column (30 m × 0.25 mm i.d. × 0.25 µm film). The GC oven program is shown in Table S6. Mass spectra were acquired in electron-impact (EI) mode at 70 eV over m/z 50–700 with an 8-min solvent delay. Data was processed in Agilent’s MassHunter Workstation 10.0. A dedicated acquisition/processing method was used for consistent retention-time alignment and peak integration. For each sterol, the most intense and unique m/z was designated the qualifier ion, with additional fragments used for confirmation when available. A signal-to-noise (S/N) threshold of 5 was used as the pass/fail criteria for all unknown sterols; only peaks exceeding this value were retained. Standard compounds ranging from 0.001 μM to 1 μM were used to construct calibration curves for quantitation of compounds. The retention time and mass spectral patterns of the following sterol standards were used as references for lipid analysis: ergosterol (Smolecule, catalog #S527372), lanosterol (Smolecule, catalog #S532452), obtusifoliol (Smolecule, catalog #S563624), zymosterol (Smolecule, catalog #S580329), 4,4-dimethyl zymosterol (Avanti catalog #700073), eburicol (Smolecule, catalog #S633611), episterol (Smolecule, catalog #S628882), and gramisterol (24-methylenelophenol or 4-methyl episterol) (Smolecule, catalog #S626191). The response for each sterol matching the standards was converted to pmol and normalized to the pmol amount of inorganic phosphate.

## ACKNOWLEDGEMENTS

We are grateful to the Selmecki laboratory and Dana Davis for helpful discussions and feedback on the manuscript. We thank Richard Bennett for sharing the RSY426 strain and Teresa O’Meara for discussions about plasmid design. This research was supported by the National Institutes of Health’s National Center for Advancing Translational Sciences, grants 1T32TR004385 and 1UM1TR004405 to EW. Funding for this work was provided by the Burroughs Wellcome Fund Investigator in the Pathogenesis of Infectious Diseases Award (#1020388) to AS, the National Institutes of Health (R01 AI143689) to AS, the University of Minnesota Graduate School Doctoral Dissertation Fellowship to NS, and in part by the National Institutes of Health (R01 AI125770) to MDP. MDP is a recipient of the Senior Research Career Scientist Award (IK6 RD001331) from the United States Department of Veterans Affairs. The content is solely the responsibility of the authors and does not necessarily represent the official views of the funding agencies.

## DATA AVAILABILITY

All whole genome sequences are available in the NCBI Sequence Read Archive repositories at BioProject Accession numbers PRJNA1338237 and PRJNA1111583. Scripts are available at https://github.com/selmeckilab/2024_Candida_lusitaniae_in_vitro_evolution.

## SUPPLEMENTAL MATERIAL

## Supplemental Tables

Table S1: Strains, Primers and Plasmids used in this study.

Table S2: Mutations detected from whole genome sequencing from *C. lusitaniae in vitro* evolution.

Table S3: Area under the curve (AUC) from growth curve +/- antifungals, MIC values, with corresponding statistics.

Table S4: Area under the curve (AUC) from growth curve +/- antifungals and +/- radicicol, with corresponding statistics.

Table S5: Median fluorescent intensity for cell wall staining with corresponding statistics.

Table S6: Abundance of sterols from targeted GC-MS with corresponding statistics.

## AUTHOR CONTRIBUTIONS

EW: Investigation, Conceptualization, Formal analysis, Methodology, Visualization, Writing original draft, Writing Review and Editing

NS: Investigation, Conceptualization, Formal analysis, Methodology, Visualization, Software development, Writing Original Draft, Writing Review and Editing

KM: Investigation, Conceptualization, Formal analysis, Methodology, Writing Original Draft,

XZ: Investigation, Conceptualization, Formal analysis, Methodology, Writing Review and Editing

DDS: Methodology, Formal analysis

NPDS: Methodology, Formal analysis

SAU: Methodology, Formal analysis

NFVDS: Methodology, Formal analysis

MDP: Resources

AS: Conceptualization, Methodology, Resources, Funding Acquisition, Writing Original Draft, Writing Review and Editing

## SUPPLEMENTARY FIGURES

**FIG. S1.**
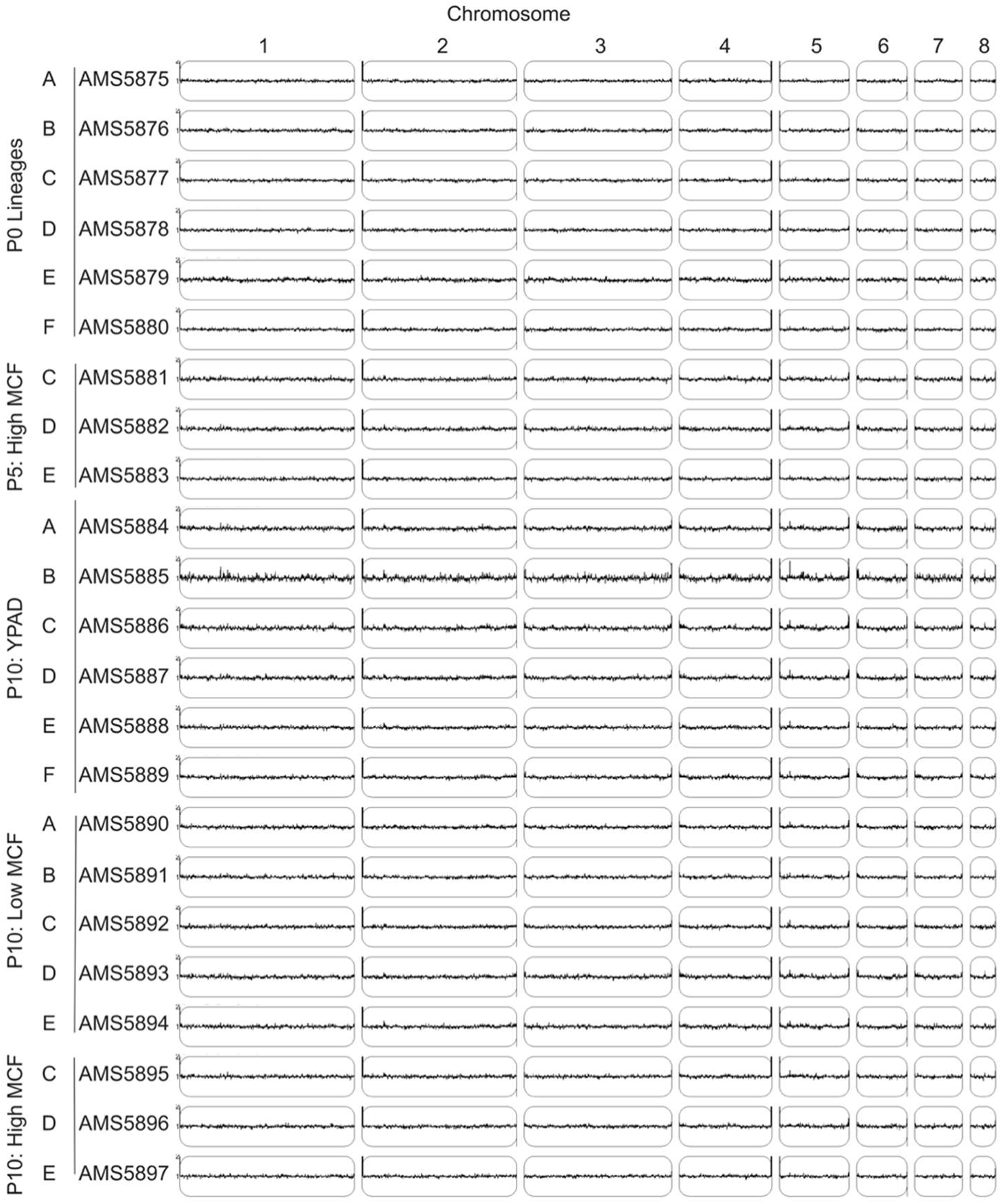
No large-scale copy number variations detected in the YPAD or micafungin-selected populations. Relative read depth (Y-axis) plotted according to genome location for all initial and evolved lineages. The *de novo* genome assembly for the drug-sensitive *C. lusitaniae* progenitor (AMS5200) was used as the reference genome and data were plotted using YMAP (140). *C. lusitaniae* is haploid and a relative copy number of 1 is expected across the genome. From top to bottom: the initial single-colonies used to start the parallel *in vitro* experiments (AMS5875-AMS5880); the “High” populations at passage 5 isolated before the increase in MCF concentration (AMS5881-AMS5883); the control YPAD populations at passage 10 (AMS5884-AMS5889); the “Low” populations at passage 10 after selection in 0.016 µg/mL MCF (AMS5890-AMS5894); and the “High” populations at passage 10 after selection in 0.032 µg/mL MCF (AMS5895-AMS5897).

**FIG S2.**
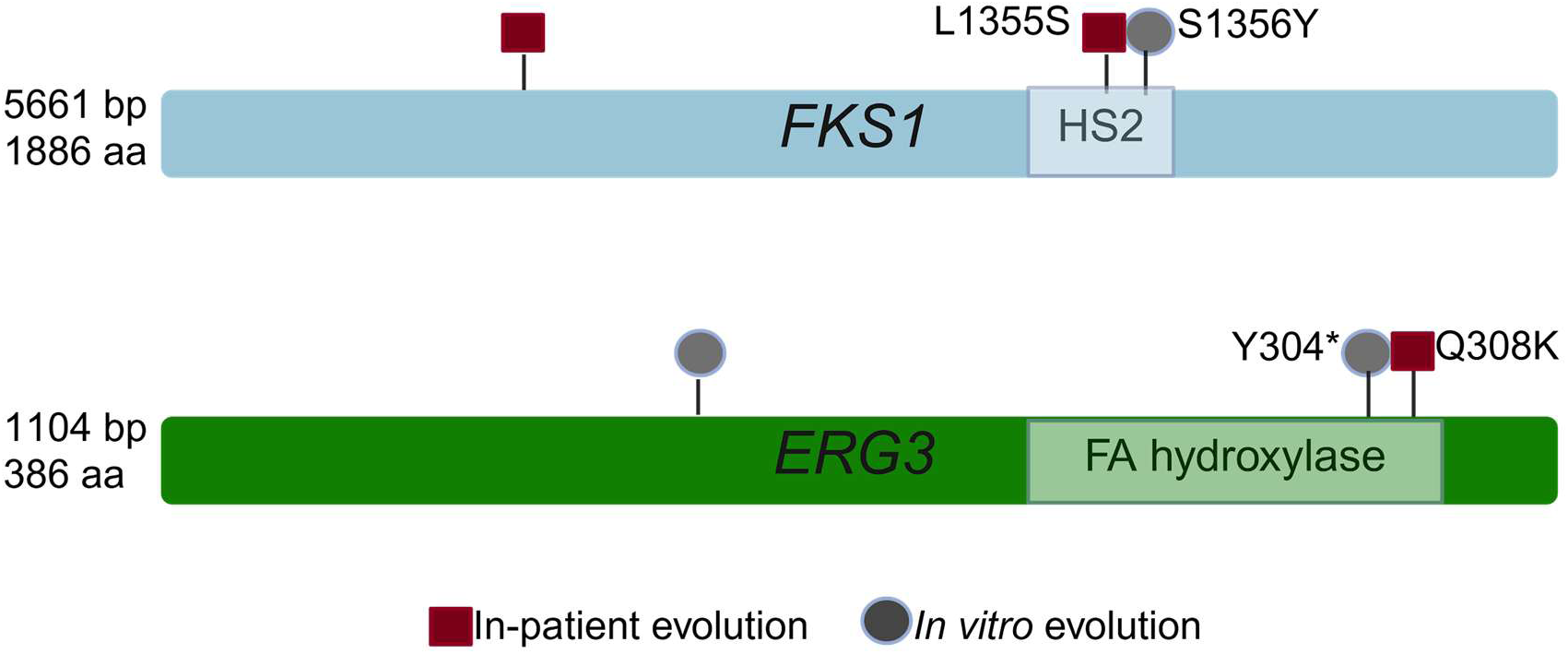
Fks1 and Erg3 domains recurrently mutated during evolution *in vitro* and within a patient receiving micafungin monotherapy. Schematic indicating the location of mutations within *FKS1* and *ERG3* across independently evolved lineages (gray circles) and the previously reported serial bloodstream isolates of *C. lusitaniae* from the same patient receiving micafungin monotherapy (red squares) (64).

**FIG. S3.**
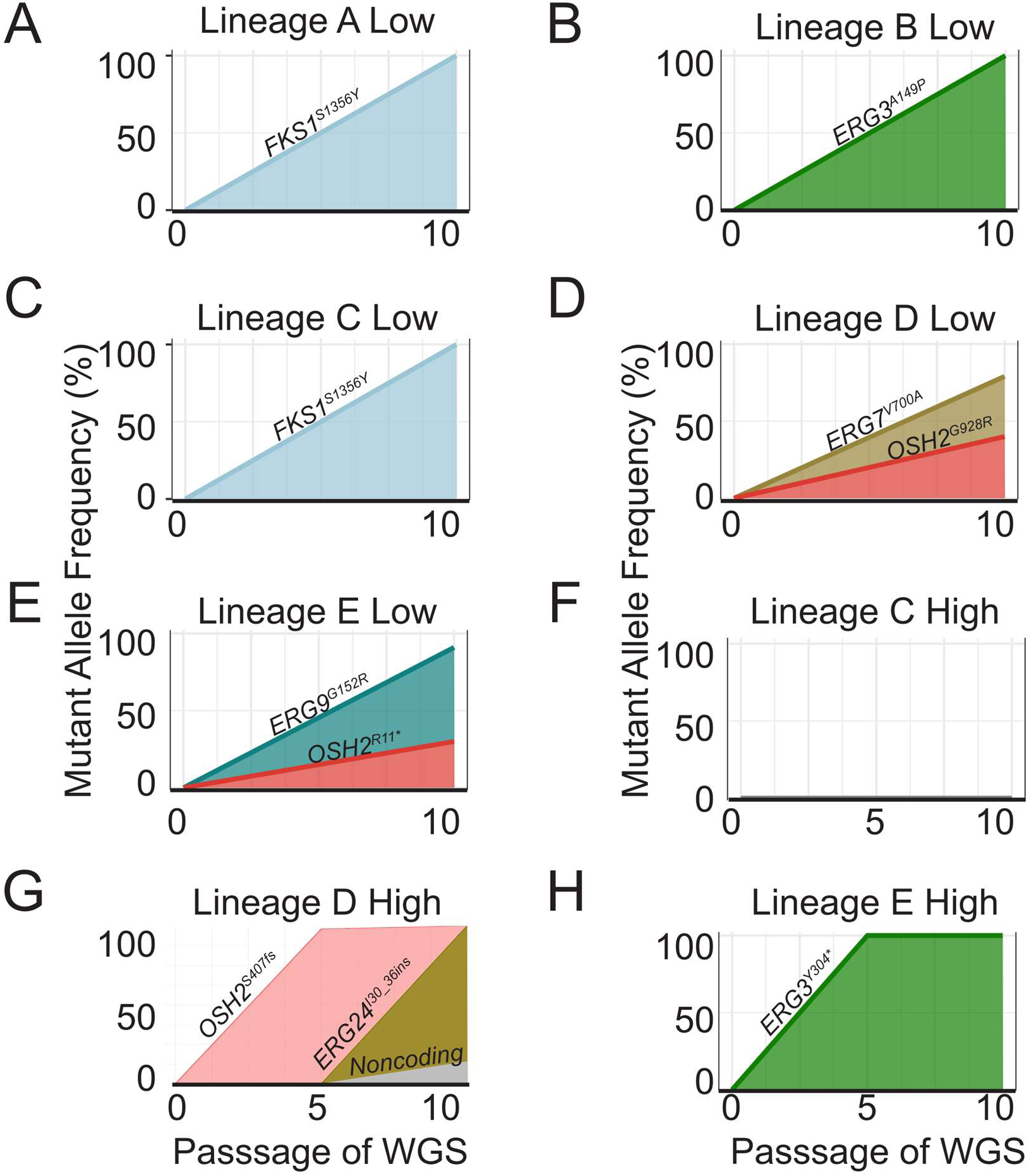
Micafungin rapidly selects for single gene variants. The mutant allele frequencies identified from population-level whole genome sequencing at passages 0 and 10 for all surviving Low MCF lineages (A-E) and for passages 0, 5, and 10 for all surviving High MCF lineages (F-H). Sequence variants are listed above colored polygon indicating frequency per population.

**FIG. S4:**
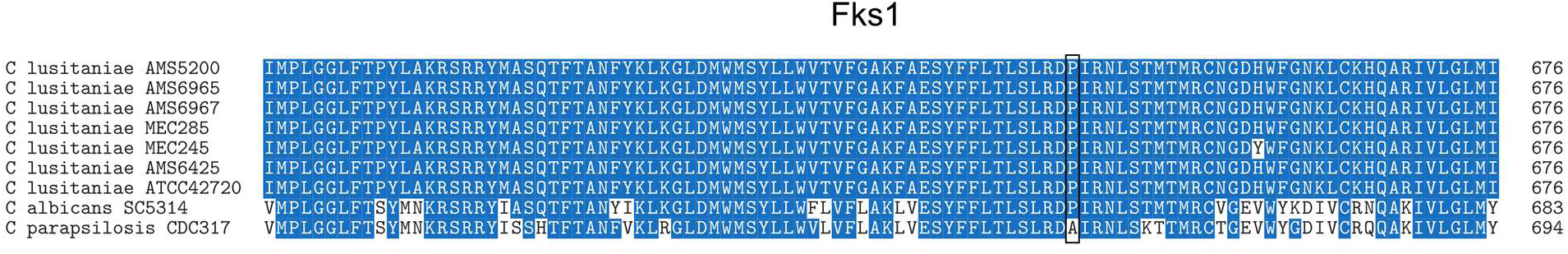
The proline-to-alanine variant in Fks1 underlying echinocandin resistance in *C. parapsilosis* is a conserved proline in *C. lusitaniae.* Multiple sequence alignment of a portion of the Fks1 protein sequence for the following 9 isolates (top to bottom): (1) *C. lusitaniae in vitro* evolved isolate with *ERG3^Y304^*\* (AMS6965, lineage B Low); (2) *C. lusitaniae* clinical isolate MEC285, from the first unrelated patient (64); (3) *C. lusitaniae in vitro* evolved isolate with *ERG3^A149P^*(AMS6967, lineage E High); (4) *C. lusitaniae* RSY426 strain (AMS6425, (87)); (5) *C. lusitaniae* clinical isolate AMS5200 and progenitor for the current study (64); (6) *C. lusitaniae* clinical isolate ATCC42720; (7) *C. lusitaniae* clinical isolate MEC245, from the second unrelated patient (64); (8) *C. albicans* SC5314 refence strain (A22) (145); (9) *C. parapsilosis* CDC317 reference strain (154). *C. parapsilosis* amino acid 660 has a unique variant (A660P, box), whereas all *C. lusitaniae* isolates have a conserved alanine residue at this position. Colors indicate conservation of amino acid, where blue is ≥ 50% conserved and white is non conserved (153).

**FIG. S5.**
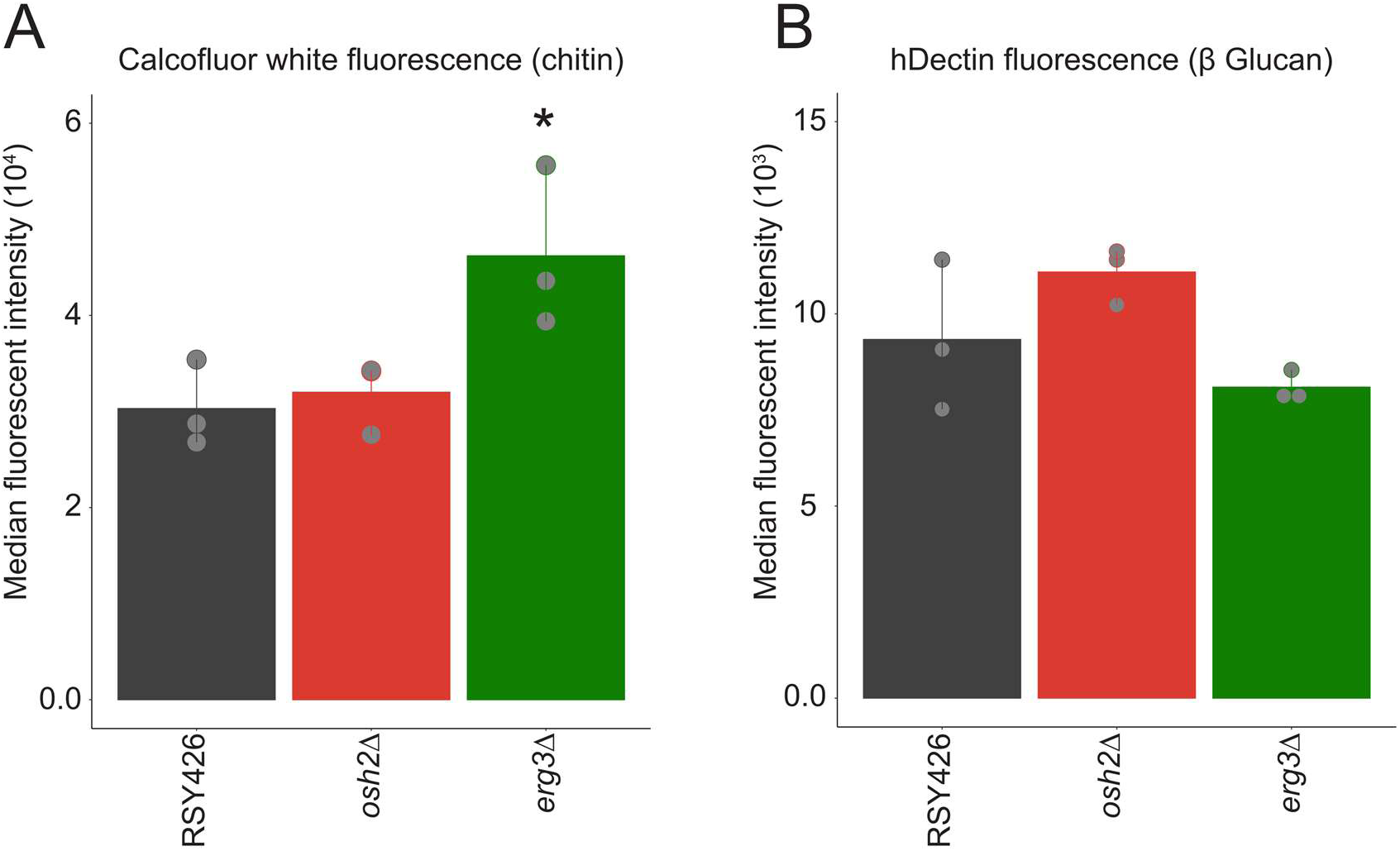
Engineered *erg3Δ* mutant has significant increase in chitin abundance. (**A**) Chitin content following staining with 10 µg/mL Calcofluor white was measured by median fluorescence intensity by flow cytometry. Asterisks indicate significant difference relative to RSY426 for the engineered mutants (*osh2Δ*, *p* = 0.94; *erg3Δ*, *p* = 0.04, ANOVA with Tukey HSD). Points indicate biological replicates; n ≥ 10,000 cells per replicate. (**B**) Median fluorescence intensity of 1,3-β-D-glucan content following staining with 15 µg/mL Fc-1-Dectin was measured by flow cytometry in for engineered mutants (*osh2Δ*, *p* = 0.99; *erg3Δ*, *p* = 0.67, ANOVA with Tukey HSD). Points indicate biological replicates; n ≥ 10,000 cells per replicate.

**FIG. S6.**
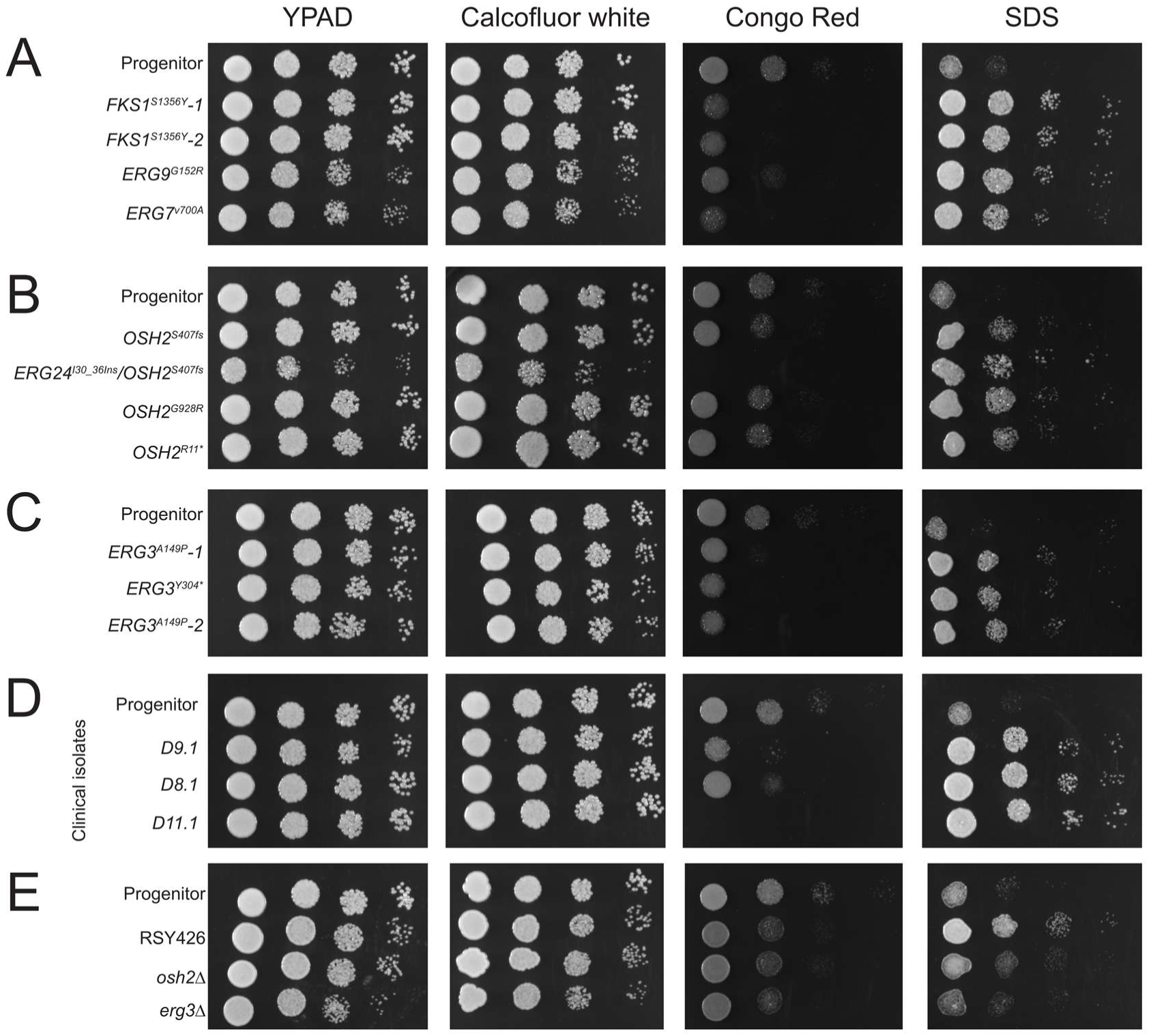
*C. lusitaniae* isolates with mutations in ergosterol biosynthesis genes have increased sensitivity to cell wall stress. Spot plate assays for *C. lustianiae* isolates on YPAD or YPAD with one of the following stresses: 15 µg/mL Calcofluor white, 150 µg/mL Congo Red, and 0.005% Sodium Dodecyl Sulfate (SDS). Serial dilutions (10^6^-10^2^) were plated. All plates were repeated in triplicate, incubated at 30°C and imaged at 24h. (**A-C**) The *C. lusitaniae* progenitor and micafungin-selected isolates. (**D**) The *C. lusitaniae* drug-sensitive progenitor and three additional clinical isolates obtained from the same patient described in (64): D9.1 (*ERG3^Q308K^*), D8.1 (*FKS1^L1355S^*), and D11.1 (*FKS1^L685F^*, *SIP3^D333E^*, *RPL8B^A161V^*, *RLM1^N181K^*). (**E**) The *C. lusitaniae* progenitor, RSY426 (*ku70::FRT*, *lig4::FRT*), and the *osh2Δ* and *erg3Δ* mutants engineered in the RSY426 background.

**FIG. S7:**
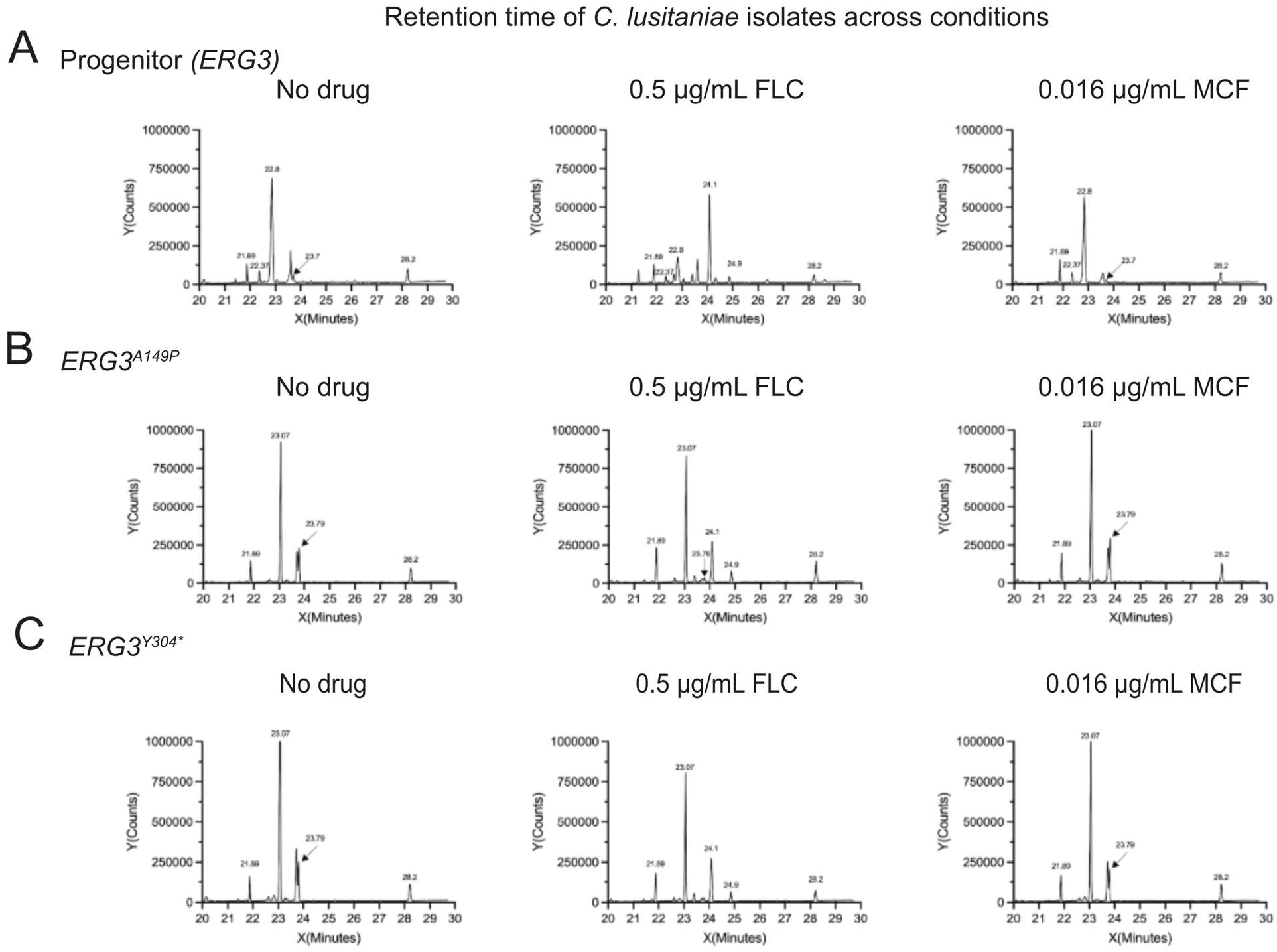
Sterol chromatograms identified from the *C. lusitaniae* progenitor and *ERG3* mutants from targeted GC-MS. Representative GC-MS chromatograms for the following isolates: (**A**) the drug-sensitive *C. lusitaniae* clinical isolate AMS5200 and progenitor for the current study (64); (**B**) the *in vitro* evolved *ERG3^A149P^*mutant; and (**C**) the *in vitro* evolved *ERG3^Y304*^* mutant. GC-MS data obtained from cells grown for 4 hours in 0 µg/mL antifungals (No drug), 0.5 µg/mL fluconazole (FLC), or 0.016 µg/mL micafungin (MCF). Retention times indicated by values adjacent to peaks for known standards.

**FIG. S8.**
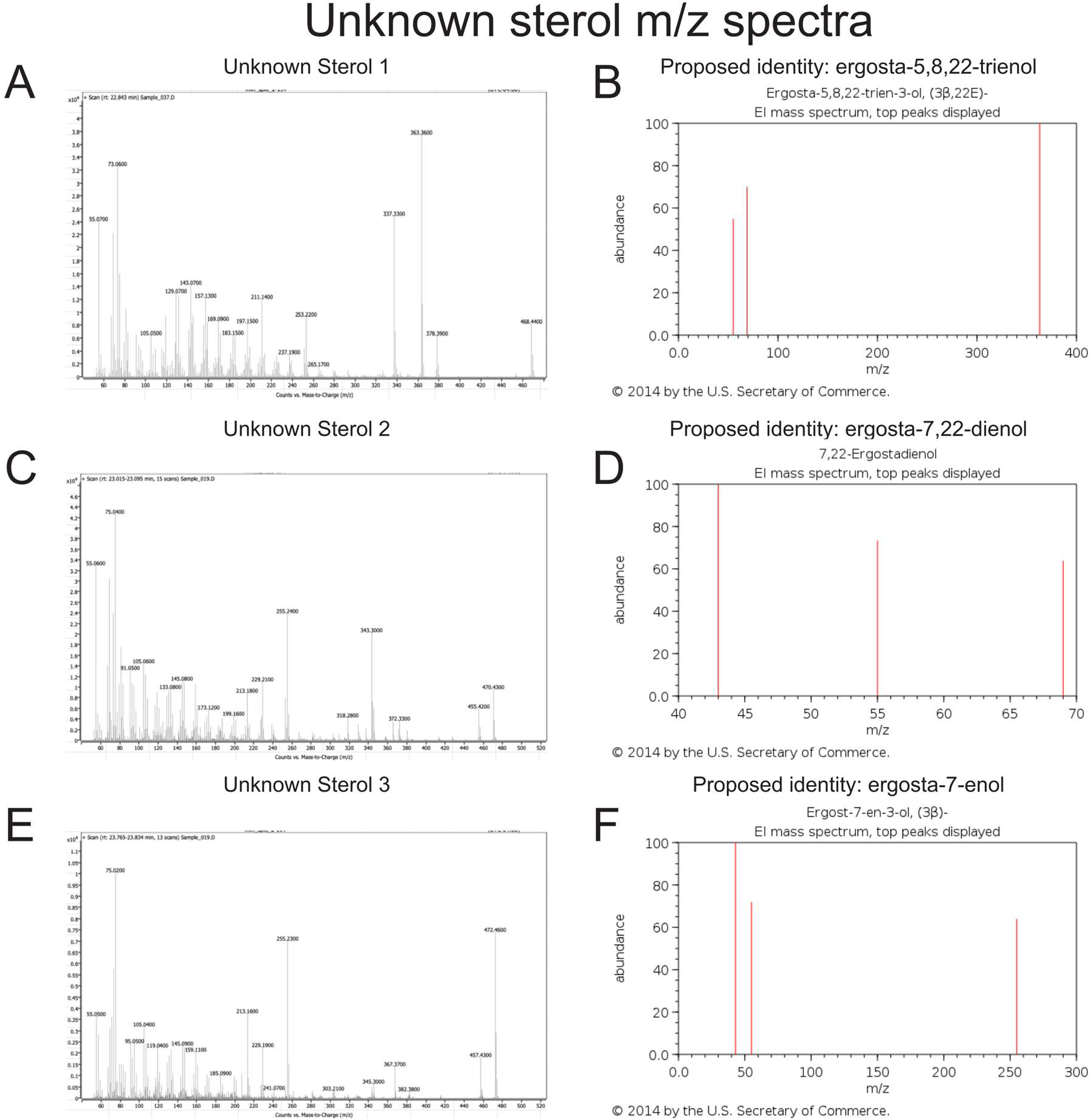
Sterols spectra from targeted GC-MS are predicted to be alternative sterol species. M/Z spectral patterns for unknown sterol species from untargeted GC-MS (labeled Unknown Sterols (**A**) 1, (**C**) 2, (**E**)3) and spectral patterns for alternative sterols predicted as unknown identity. Alternative sterol spectra adapted from PubChem (ID: (**B**) 70147717, (**D**) 5283628, (**F**) 91691464).

